# Major surgery induces acute changes in DNA methylation associated with activation of the immune response

**DOI:** 10.1101/706184

**Authors:** Ryoichi Sadahiro, Bridget Knight, Ffion James, Eilis Hannon, John Charity, Ian R. Daniels, Joe Burrage, Olivia Knox, Bethany Crawford, Neil J. Smart, Jonathan Mill

## Abstract

**Background:** Surgery is an invasive procedure evoking acute inflammatory and immune responses that are believed to mediate risk for postoperative complications including cognitive dysfunction and delirium. Although the specific mechanisms driving these responses have not been well-characterized, they are hypothesized to involve the epigenetic regulation of gene expression. We quantified genome-wide levels of DNA methylation in purified peripheral blood mononuclear cells (PBMCs) longitudinally collected from 55 elderly patients undergoing three types of major surgery (elective colorectal and hip replacement surgery, and emergency hip fracture surgery), comparing samples collected at baseline to those collected immediately post-operatively and at discharge from hospital.

**Results:** Major surgery was associated with acute changes in DNA methylation at sites annotated to immune system genes, paralleling changes in serum-levels of markers including C-reactive protein (CRP) and Interleukin 6 (IL-6) measured in the same individuals. Although many of the observed changes in DNA methylation are consistent across the three types of surgery, there is notable heterogeneity between surgery types at certain loci. The acute changes in DNA methylation induced by surgery are relatively stable in the postoperative period, generally persisting until discharge from hospital.

**Conclusions:** Our results highlight the dramatic alterations in gene regulation induced by invasive surgery, primarily reflecting upregulation of the immune system in response to trauma, wound healing and anaesthesia.

## Background

Surgery is an invasive procedure that elicits dramatic challenges to normal regulatory and homeostatic processes in the body. Importantly, surgery evokes acute inflammatory and immunological responses, which results directly from the trauma of surgery itself that act to modulate wound healing and other physiological pathways (1–3). Of note, changes to the immunological milieu following major surgery are thought to mediate risk for a number of postoperative complications including nosocomial infections (4), and postoperative cognitive dysfunction and delirium (5).

Post-operatively the immune system acts to promote tissue regeneration by influencing signal transduction, cell migration, differentiation, proliferation, and tissue remodelling (6, 7). Although the specific mechanisms driving these responses have not been well-characterized, they are hypothesized to involve the epigenetic regulation of gene expression (8). Epigenetic processes act to dynamically control transcription via modifications to DNA, histone proteins, and chromatin acting independently of DNA sequence variation, and are known to regulate the function of immune cells during health and disease (9). Of note, changes in DNA methylation have been reported to mediate postsurgical pain sensitivity (10) and response to anaesthesia (11, 12) and opioids (13, 14).

In this study we explored the hypothesis that immunological modulation following surgery is associated with acute changes in DNA methylation. We longitudinally profiled DNA methylation in purified peripheral blood mononuclear cells (PBMCs) collected from elderly patients undergoing different types of major surgery at three time-points. Our study identifies dramatic shifts in DNA methylation in the vicinity of genes involved in immune regulation, with these changes being relatively stable across the post-operative period.

## Results

### Methodological overview

Using a longitudinal study design, we quantified DNA methylation across the genome in a cohort of 30 elderly patients (average age = 77.9 years (range 62-91)) undergoing major surgery (either elective colorectal or hip replacement surgery, and emergency hip surgery following fracture). From each individual, detailed blood measures were collected and DNA methylation was profiled using the Illumina Human Methylation 450 array (“450K array”) (Illumina Inc. San Diego, California) in DNA samples isolated from peripheral blood mononuclear cells (PBMCs) collected at three time-points: i) immediately before surgery (baseline), ii) in the morning of post-operative day 1 (POD1) and iii) between post-operative day 4 and 7 (POD4/7) prior to discharge from hospital. Our analyses focused on identifying differentially methylated positions (DMPs) and regions (DMRs) reflecting acute changes induced by major surgery (see **Materials and Methods**). An overview of the study design and sampling strategy is given (**Supplementary Figure 1)** and an overview of the patients included in this study is presented (**Table 1)**.

**Table 1.** An overview of the surgical patients enrolled in this study. ADL = activity of daily life. ASA = American Society of Anaesthesiologists.

| **Variables** | **All** | **Colorectal** | **Hip Elective** | **Hip Fracture** | ***P*** |
| --- | --- | --- | --- | --- | --- |
| Number of Subjects | 30 | 11 | 10 | 9 | - |
| Age | 77.9±6.4 | 73.6±5.8 | 79.4±4.6 | 81.3±6.4 | 1.24E-02 |
| Sex (M :F ) | 13 : 17 | 7 : 4 | 4 : 6 | 2 : 7 | - |
| Smoking (Never : Ex : Current) | 21 : 9 : 0 | 8 : 3 : 0 | 5 : 5 : 0 | 7 : 2 : 0 | - |
| Katz index of ADL | 11.9±0.5 | 12±0 | 12±0 | 11.7±0.9 | 3.22E-01 |
| Charlson's comorbidity score | 1.7±1.3 | 2.0±1.3 | 1.9±1.2 | 1.2±1.3 | 3.69E-01 |
| ASA class (1-2 : 3-4) | 23 : 7 | 9 : 2 | 8 : 2 | 6 : 3 | - |
| Surgery time (hr) | 2.2±1.1 | 3.0±1.0 | 1.4±0.5 | 2.0±1.1 | 1.13E-02 |
| Anaesthesia time (hr) | 2.9±1.1 | 3.7±1.1 | 2.3±0.7 | 2.8±1.1 | 1.62E-03 |
| Time to POD1 sample (hr) | 18.1±3.2 | 17.4±3.3 | 20.0±2.6 | 16.9±3.0 | 6.35E-02 |
| Time to POD4/7 sample (days) | 4.8±1.0 | 5.5±1.2 | 4.2±0.4 | 4.6±0.7 | 4.24E-03 |

### Major surgery is associated with acute changes in DNA methylation, primarily at sites annotated to immune system genes

We first examined acute changes in DNA methylation following major surgery, comparing baseline PBMC samples to those at POD1, collected an average of 18.1±3.2 hours after surgery. In total, we identified 88 DMPs passing an experiment-wide significance threshold (P < 2.0E-07), with 5,655 DMPs passing a more relaxed *“discovery”* significance threshold (P < 5.0E-05) that was used to select genes for subsequent gene ontology (GO) pathway analyses (**Supplementary Figure 2** and **Supplementary Table 1**). The top ranked DMP (cg15412772), which was significantly hypomethylated following major surgery (DNA methylation change = −4.23%, P = 8.08E-10), is located on chromosome 17 approximately 4kb downstream of *CSF3*, which encodes a granulocyte colony-stimulating factor that acts to stimulate granulopoiesis and works against neutropenia (15). We used *comb-p* to identify spatially-correlated regions of differential DNA methylation, identifying 913 surgery-associated DMRs (**Supplementary Table 2**) (Sidak-corrected P < 0.05) (16). The top-ranked DMR spans 1,212 bp, incorporating 19 DNA methylation sites on chromosome 6 (P = 2.58E-26), overlapping the transcription start site (TSS) of the gene encoding lymphotoxin-A (*LTA*), which has an important role in regulating the immune system (17). GO term enrichment analysis on genes annotated to the *“discovery”* DMPs highlighted a highly-significant enrichment for functional pathways associated with immune function amongst surgery-induced DMPs, including categories such as *“immune system process”* (P = 9.51E-24), *“response to wounding”* (P = 1.17E-19) and *“positive regulation of immune response”* (P = 4.50E-12) (**Supplementary Table 3**). To explore further the acute immune response following major surgery we measured blood serum levels of C-reactive protein (CRP) and Interleukin 6 (IL-6) in the same samples; CRP is an acute phase protein that rises in response to inflammation through hepatic synthesis in response to IL-6, a pro-inflammatory cytokine that is synthesised at the site of injury (18, 19). Compared to baseline we observed a highly significant increase in both serum CRP (baseline: median CRP = 6.0 mg/ml; POD1: median CRP = 65 mg/ml, P = 4.58E-03) (**Supplementary Table 4** and **Figure 1A**) and IL-6 (baseline: median IL-6 = 6.7 pg/ml; POD1: median IL-6 = 114.3 pg/ml, P = 7.92E-06) (**Supplementary Table 4** and **Figure 1B**) at POD1.

**Figure 1 -.**
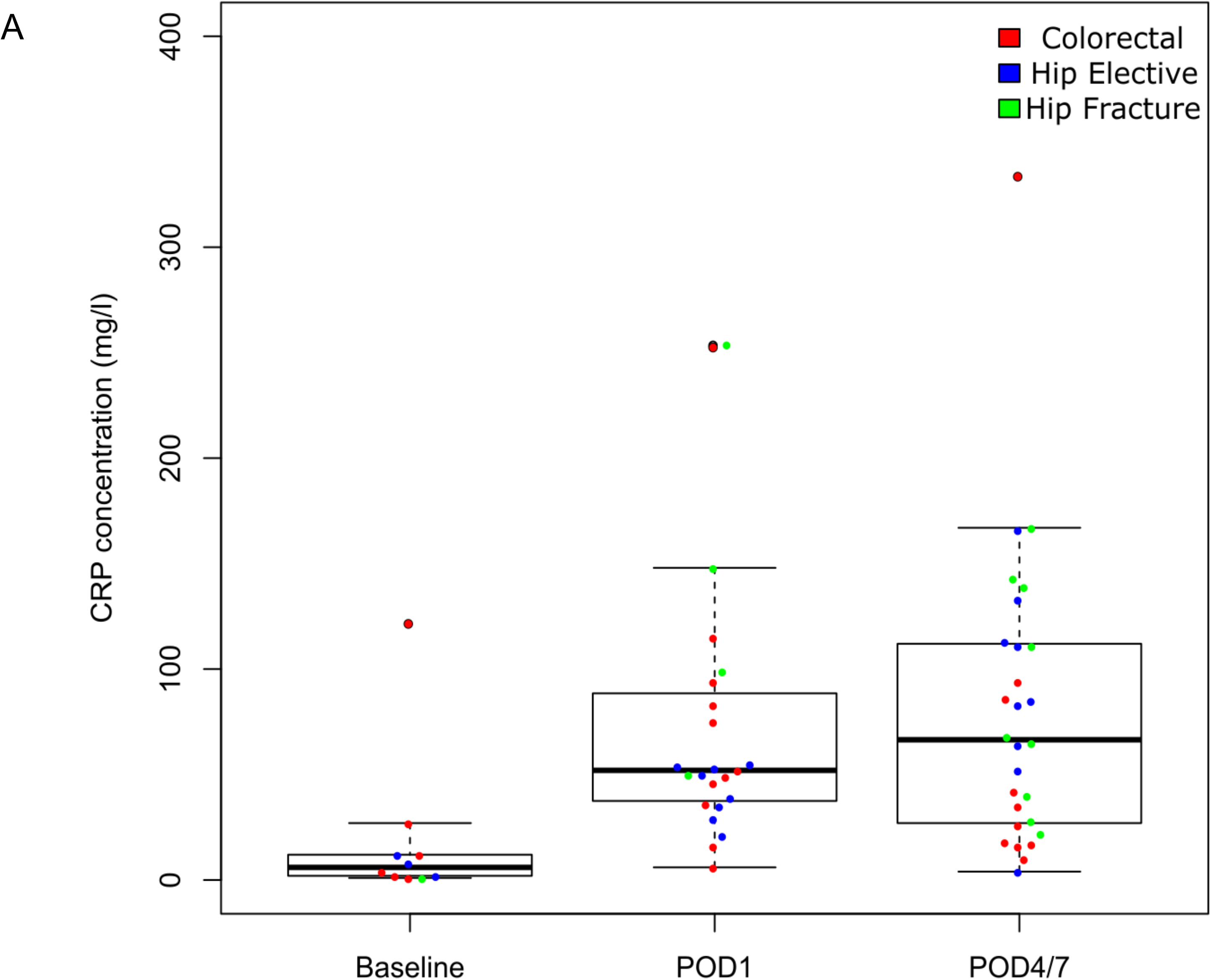
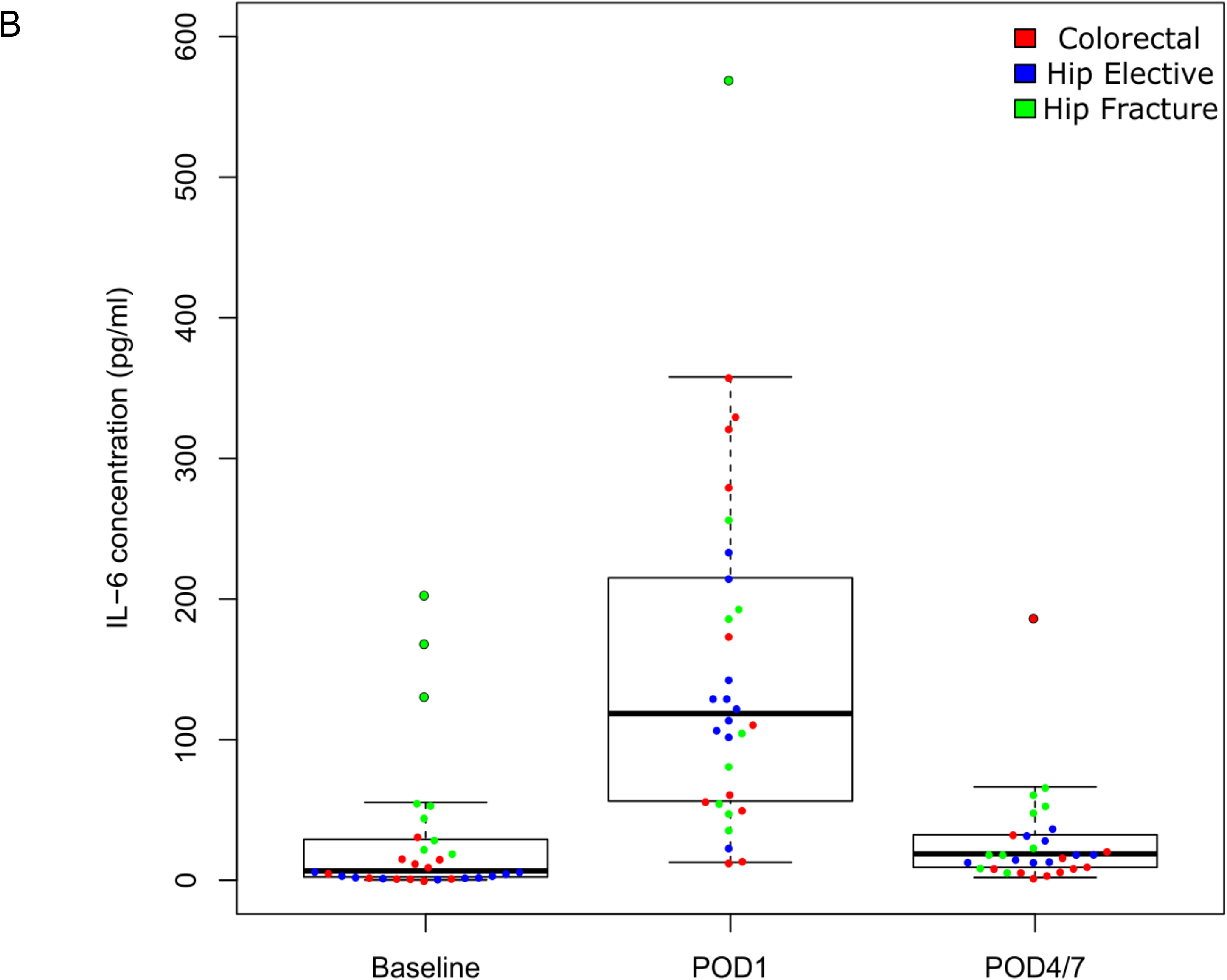
Surgery is associated with acute increases in markers of inflammation. Shown are levels of serum C-reactive protein (CRP) and Interleukin-6 (IL-6) at baseline and after surgery. Both serum CRP **(A)** and IL-6 **(B)** are significantly elevated at POD1 following major surgery (CRP; P = 4.58E-03, IL-6; P = 1.37E-05). Colors denote surgery type: colorectal surgery = red, hip elective surgery = blue, and hip fracture surgery = green.

### A subset of surgery-associated changes in DNA methylation are independent of blood cell heterogeneity

Although our DNA methylation analyses were performed on purified PBMC samples, given the highly-significant enrichment of surgery-associated DMPs in the vicinity of genes involved in immune function and cell proliferation, the dramatic surgery-induced changes in CRP and IL-6, and the large observed differences in blood cell proportions following surgery (**Supplementary Figure 4** and **Supplementary Table 4**), we hypothesized that it is likely to be important to further control for variable cell populations in our DNA methylation analysis of PBMC samples. We used an established a method to derive cell proportion estimates from our PBMC DNA methylation data (see **Materials and Methods**), identifying significant acute increases following surgery (between baseline and POD1) in estimates of plasmablasts (P = 2 28E-02) and monocytes (P = 4.43E-05), and significant decreases in naïve CD8 T cells (P = 4.96E-02), CD8 T cells (P = 1.22E-02), CD4 T cells (P = 1.19E-03), and natural killer cells (P = 5.97E-03) (see **Supplementary Table 5**) (20). As expected, the estimated level of granulocytes in our isolated PBMCs was very low (mean (±SD) granulocyte proportion = 0.0129 (±0.028)). For the majority of samples we were able to also empirically measure actual blood cell counts; although the specific cell-types derived from DNA methylation data do not overlap entirely with those measured in whole blood, we observed a highly significant correlation between empirically-measured and estimated ratio of lymphocytes and monocytes (r = 0.713, P = 8.53E-15, **Supplementary Figure 5**), indicating that the derived cell proportion estimates are likely to reflect actual levels in our samples. We therefore used a multi-level regression model including the estimated proportions of plasmablasts, CD8+CD28-CD45RA-T cells, naive CD8 T cells, CD4T cells, natural killer cells and monocytes for each sample (derived from the DNA methylation data), in addition to age and sex, to further explore acute surgery-associated DMPs between baseline and POD1 (see **Materials and Methods**). Although there was a strong correlation for both effect sizes (r = 0.955, P = 2.20E-47, **Supplementary Figure 6**) and P values (r = 0.621, P = 6.45E-11, **Supplementary Figure 7**) between models for DMPs identified in our uncorrected model, the cell-proportion-corrected model identified a smaller number of significant surgery-associated changes. In total, we identified 11 DMPs passing an experiment-wide significance threshold P < 2.0E-07 (**Figure 2A** and **Table 2**) and 475 DMPs passing a more relaxed *“discovery”* significance threshold (P < 5.0E-05) (**Supplementary Figure 8** and **Supplementary Table 6**). The top ranked DMP (cg26022992), which was significantly hypermethylated following major surgery (DNA methylation change = −2.04%, P = 1.02E-08), is located on chromosome 1 in the 5’ untranslated region of *RBM8A*, a gene encoding a protein with a conserved RNA-binding motif that is involved in splicing. We again used *comb-p* (16) to identify spatially-correlated regions of differential DNA methylation, with the top-ranked DMR spanning nine hypomethylated sites on chromosome 7 immediately upstream of *PON3*, a member of the paraoxonase family that associates with high-density lipoprotein and has anti-inflammatory and anti-oxidant properties (21) (Sidak-corrected P = 1.43E-4) (**Figure 3A** and **Supplementary Table 7**). GO term enrichment analysis of genes annotated to the list of ‘*discovery*’ DMPs again highlighted a significant enrichment for functional pathways associated with immune function amongst surgery-induced DMPs, for example *“response to interleukin-15”* (P = 3.44E-10), in addition to some more physiological pathways including *“regulation of muscle adaptation”* (P = 5.23E-06) and *“respiratory system process”* (P = 5.91E-05) (**Supplementary Table 8**).

**Figure 2.**
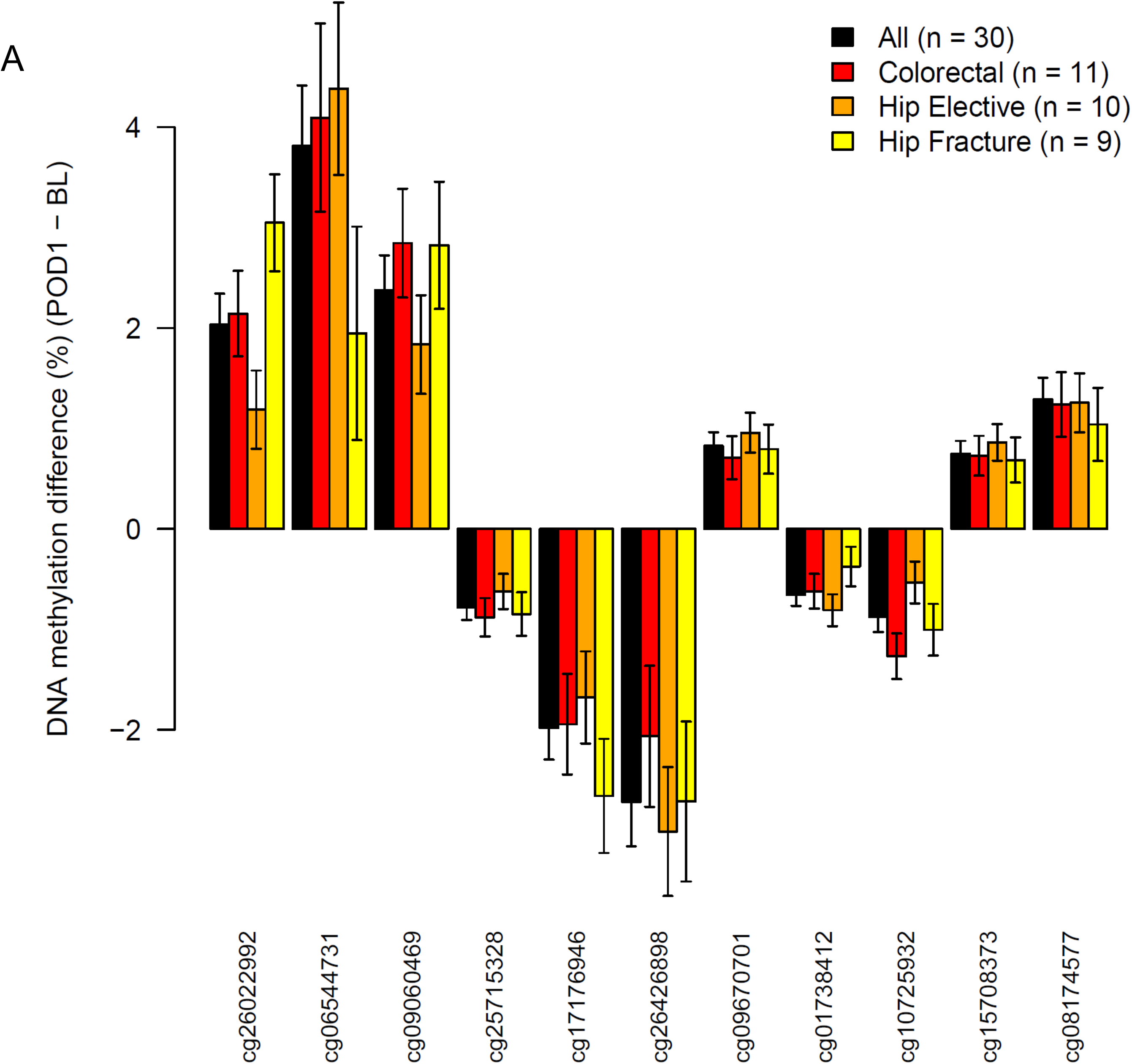
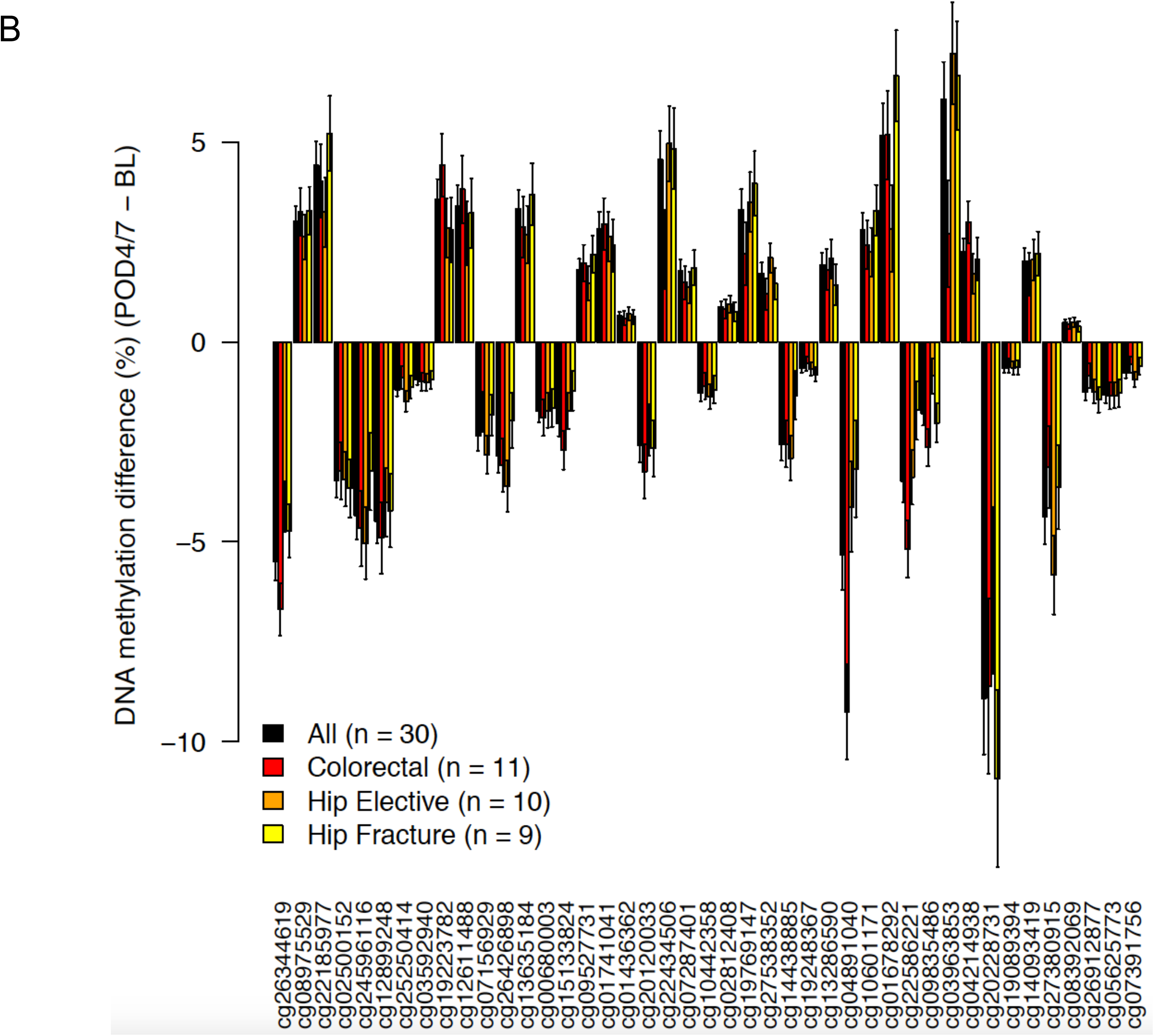
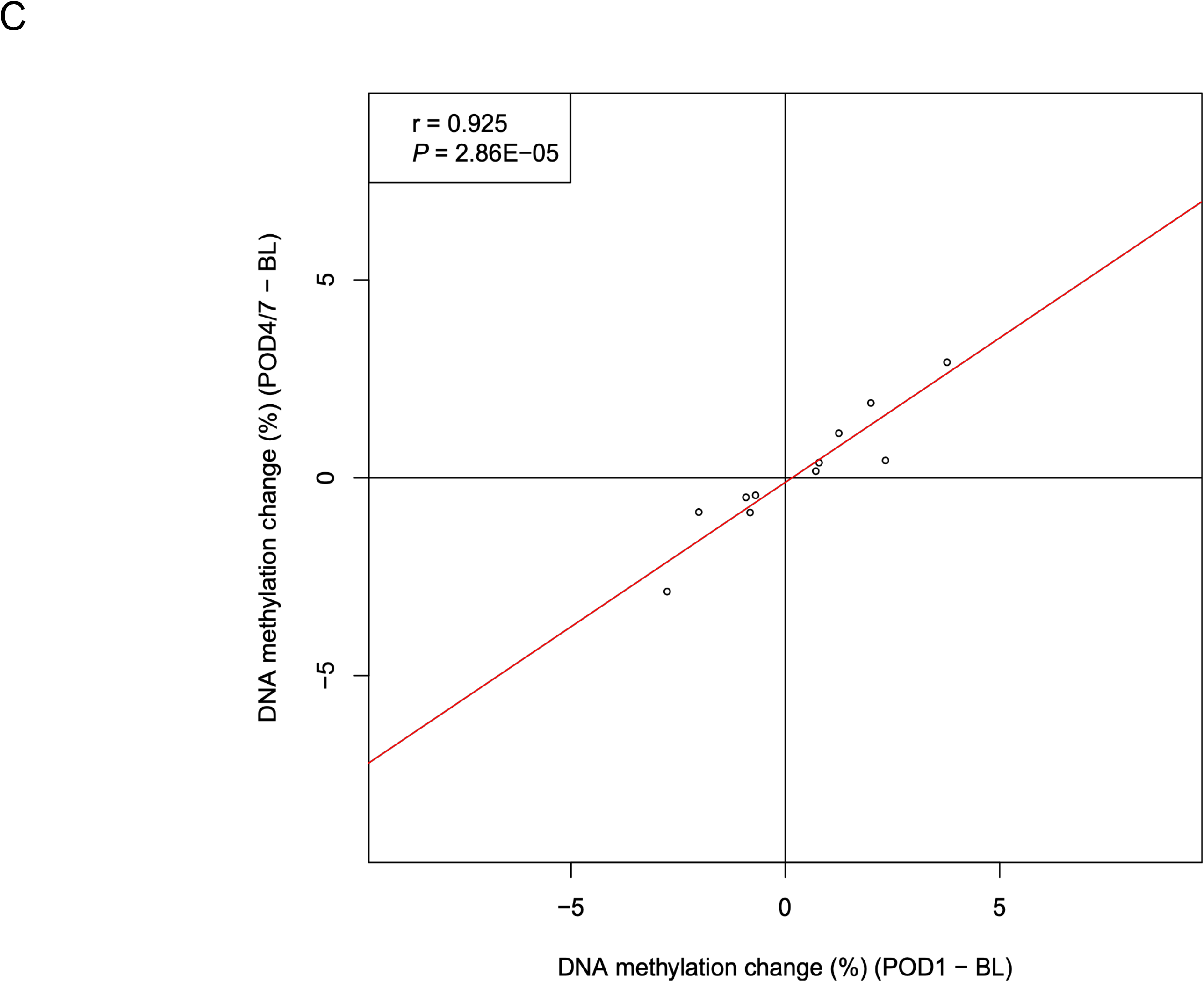
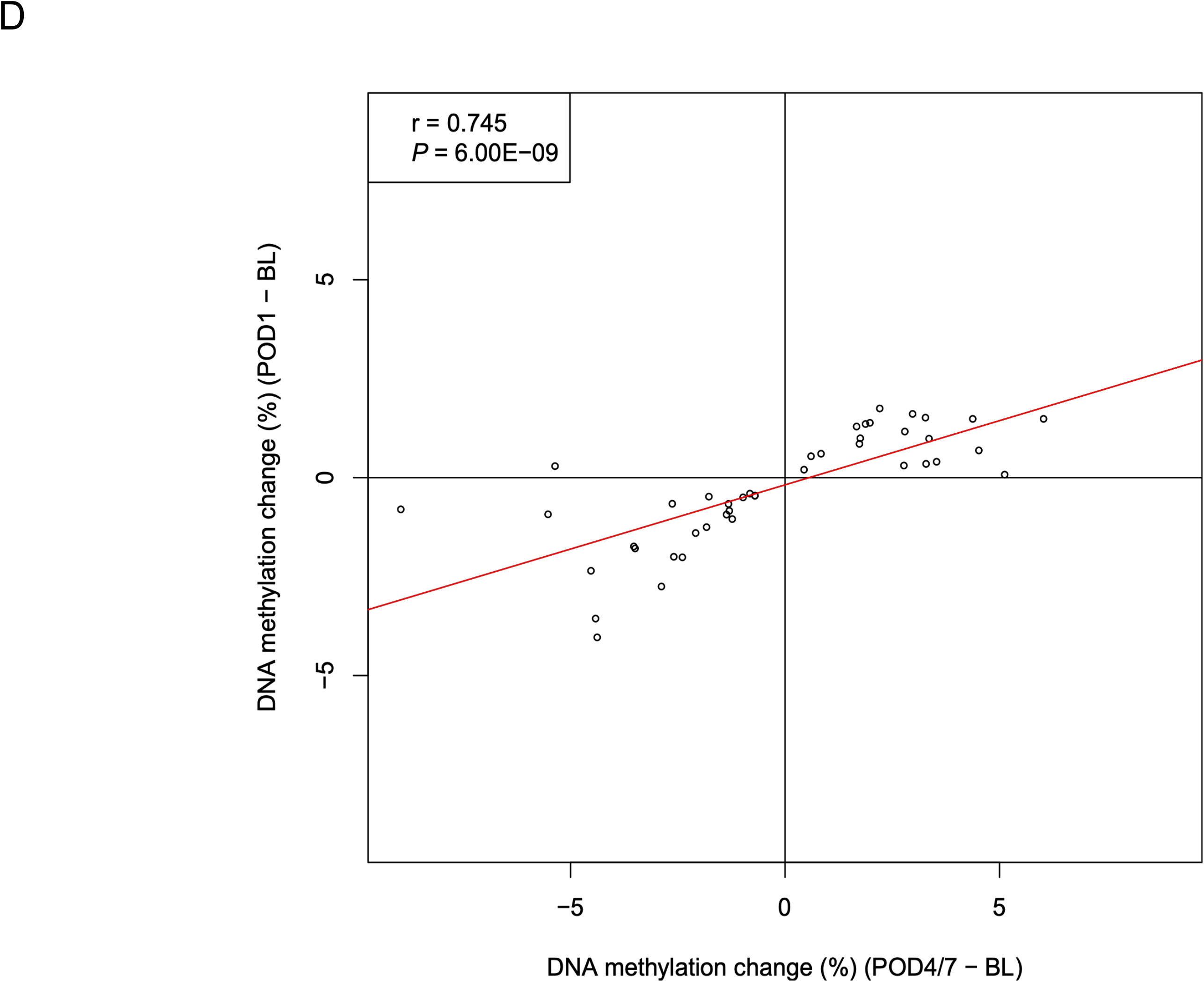
Differentially methylated positions associated with major surgery. **A)** 11 DMPs passed our experiment-wide significance threshold at POD1 (P < 2.0E-07). Results for each of these sites is given in Table 2. **B)** 43 DMPs passed our experiment-wide significance threshold at POD4/7 (P < 2.0E-07). Results for each of these sites is given in Table 3. There was a strong correlation of changes observed at POD1 and POD4/7 for DMPs identified at both time-points: **C)** POD1 DMPs: corr = 0.925, P = 2.86E-05; D) POD4/7 DMPs: corr = 0.745, P = 6.00E-09.

**Table 2.**
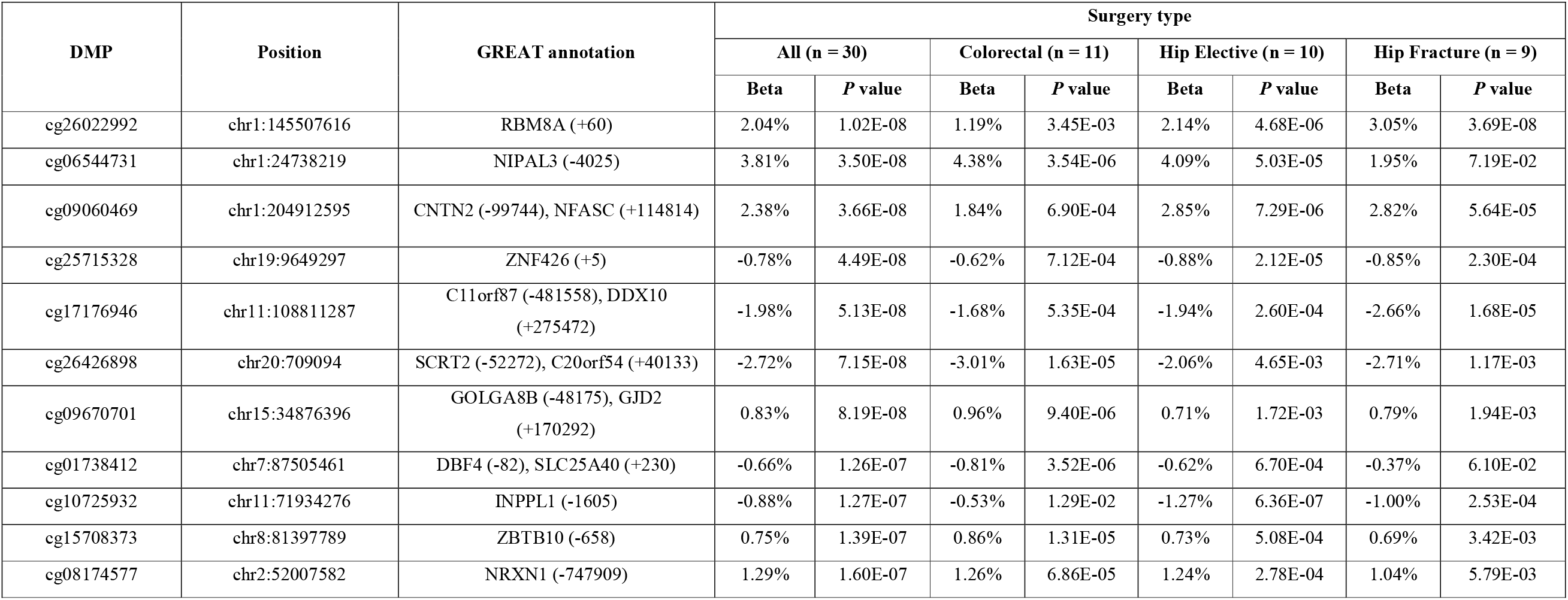
Sites characterized by significant changes in DNA methylation on post-operative day 1 compared to baseline. Shown for each DMP are the genomic location (hg19) and genic annotation derived using GREAT (35). Association statistics are presented across all patients, and separately for each surgery type.

| DMP | Position | GREAT annotation | Surgery type |  |  |  |  |  |  |  |
| --- | --- | --- | --- | --- | --- | --- | --- | --- | --- | --- |
|  |  |  | All (n = 30) |  | Colorectal (n = 11) |  | Hip Elective (n = 10) |  | Hip Fracture (n = 9) |  |
|  |  |  | Beta | P value | Beta | P value | Beta | P value | Beta | P value |
| cg26022992 | chr1:145507616 | RBM8A (+60) | 2.04% | 1.02E-08 | 1.19% | 3.45E-03 | 2.14% | 4.68E-06 | 3.05% | 3.69E-08 |
| cg06544731 | chr1:24738219 | NIPAL3 (-4025) | 3.81% | 3.50E-08 | 4.38% | 3.54E-06 | 4.09% | 5.03E-05 | 1.95% | 7.19E-02 |
| cg09060469 | chr1:204912595 | CNTN2 (-99744), NFASC (+114814) | 2.38% | 3.66E-08 | 1.84% | 6.90E-04 | 2.85% | 7.29E-06 | 2.82% | 5.64E-05 |
| cg25715328 | chr19:9649297 | ZNF426 (+5) | -0.78% | 4.49E-08 | -0.62% | 7.12E-04 | -0.88% | 2.12E-05 | -0.85% | 2.30E-04 |
| cg17176946 | chr11:108811287 | C11orf87 (-481558), DDX10 (+275472) | -1.98% | 5.13E-08 | -1.68% | 5.35E-04 | -1.94% | 2.60E-04 | -2.66% | 1.68E-05 |
| cg26426898 | chr20:709094 | SCRT2 (-52272), C20orf54 (+40133) | -2.72% | 7.15E-08 | -3.01% | 1.63E-05 | -2.06% | 4.65E-03 | -2.71% | 1.17E-03 |
| cg09670701 | chr15:34876396 | GOLGA8B (-48175), GJD2 (+170292) | 0.83% | 8.19E-08 | 0.96% | 9.40E-06 | 0.71% | 1.72E-03 | 0.79% | 1.94E-03 |
| cg01738412 | chr7:87505461 | DBF4 (-82), SLC25A40 (+230) | -0.66% | 1.26E-07 | -0.81% | 3.52E-06 | -0.62% | 6.70E-04 | -0.37% | 6.10E-02 |
| cg10725932 | chr11:71934276 | INPPL1 (-1605) | -0.88% | 1.27E-07 | -0.53% | 1.29E-02 | -1.27% | 6.36E-07 | -1.00% | 2.53E-04 |
| cg15708373 | chr8:81397789 | ZBTB10 (-658) | 0.75% | 1.39E-07 | 0.86% | 1.31E-05 | 0.73% | 5.08E-04 | 0.69% | 3.42E-03 |
| cg08174577 | chr2:52007582 | NRXN1 (-747909) | 1.29% | 1.60E-07 | 1.26% | 6.86E-05 | 1.24% | 2.78E-04 | 1.04% | 5.79E-03 |

**Figure 3.**
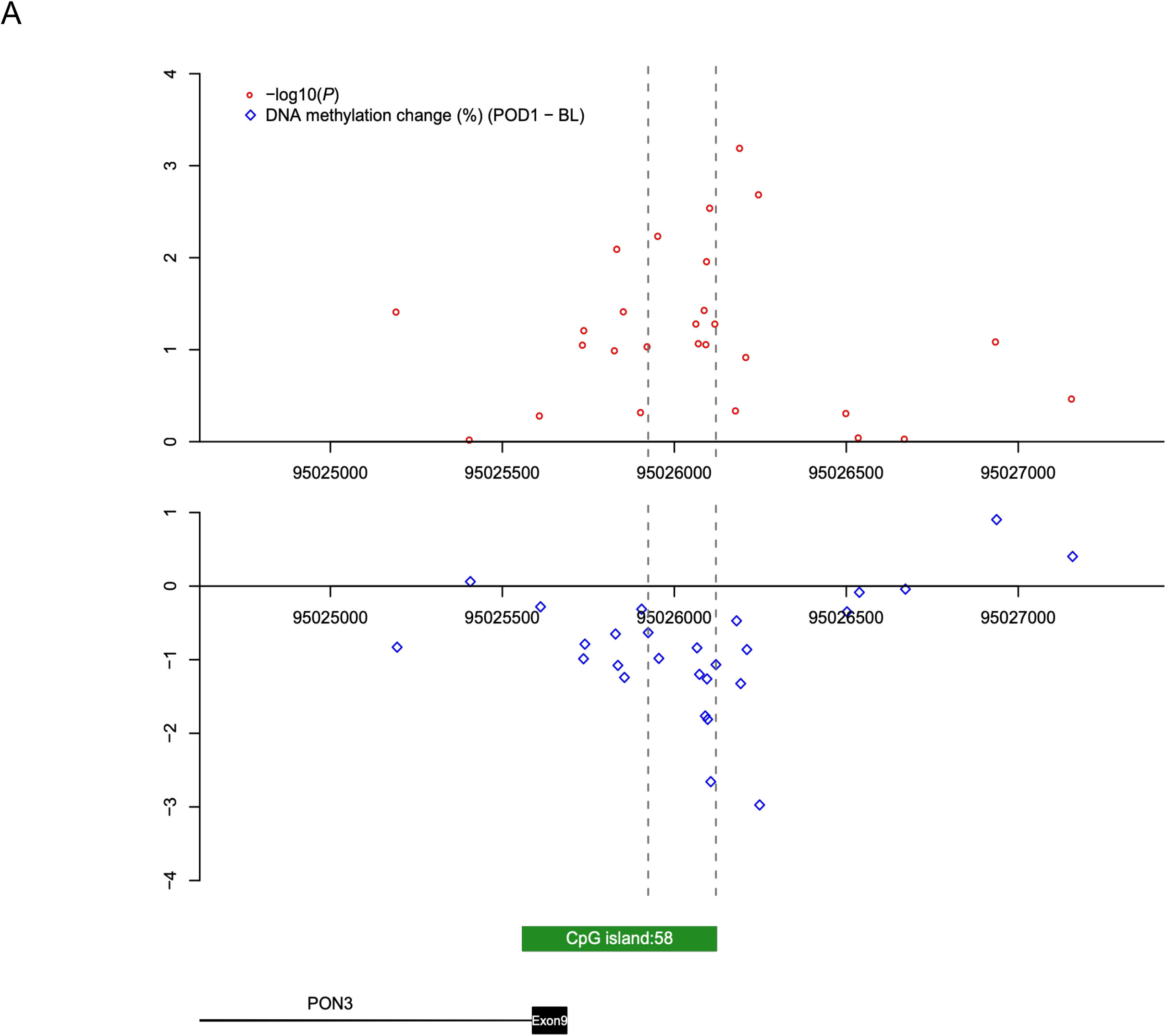
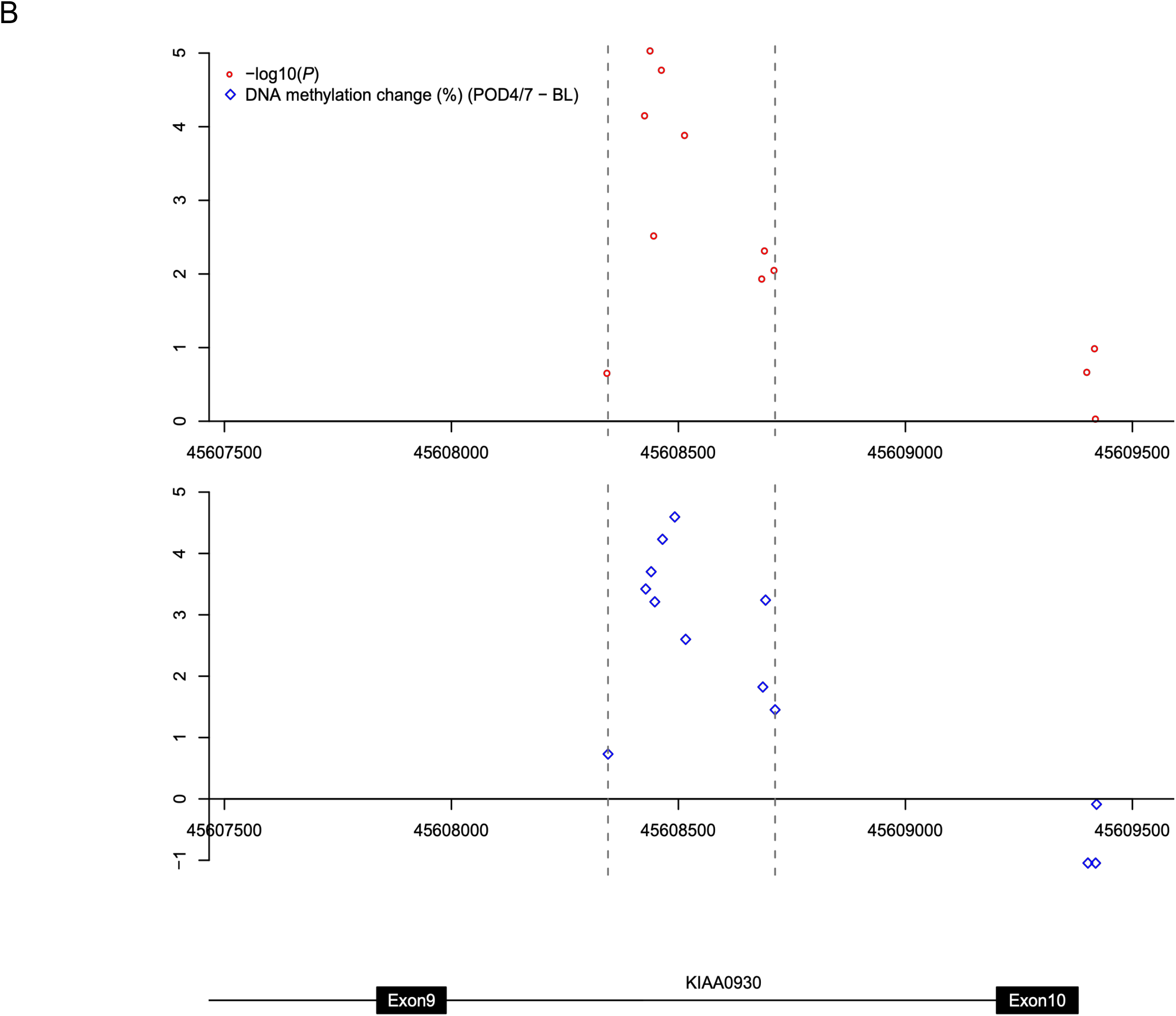
Differentially methylated regions associated with major surgery. **A)** A DMR on chromosome 7 annotated to the promoter region of PON3 which was hypomethylated (Sidak-corrected P = 1.43E-04) at POD1. **B)** A DMR on chromosome 22 within KIAA0930 which was hypermethylated (Sidak-corrected P = 3.39E-17) at POD4/7. −log10(P) values and effect sizes (%, average DNA methylation difference) are shown as red circles and blue diamonds, respectively.

### Surgery-associated DNA methylation differences are relatively stable until discharge from hospital

We next extended our analysis to characterize more long-term changes in DNA methylation following major surgery (i.e. those occurring between BL and POD4/7), exploring the stability of acute surgery-induced changes identified at POD1. In total we identified 43 DMPs characterized by an experiment-wide significant difference (P < 2.0E-07) at POD4/7 compared to BL using a model correcting for derived blood cell-type proportions (**Figure 2B** and **Table 3**), with 909 DMPs identified at a more relaxed “discovery” significance threshold (P < 5.0E-05) (**Supplementary Figure 9** and **Supplementary Table 9**). The top-ranked DMP at POD4/7 (cg26344619), which is significantly hypomethylated following surgery (DNA methylation change = −5.49%, P = 1.13E-13), is located within the first intron of *FLVCR2* on chromosome 14, a gene encoding a transmembrane protein involved in heme and calcium transport (22, 23). The top-ranked DMR at POD4/7 spans ten hypermethylated sites on chromosome 22 in the promoter region of *KIAA0930* (Sidak-corrected P = 3.39E-17) (**Figure 3B** and **Supplementary Table 7**). Functional pathways enriched amongst genes annotated to DMPs at POD4/7 again included immune response pathways such as “cellular response to interleukin-6” (P = 3.31E-06) in addition to several cytoskeletal pathways including “actomyosin” (P = 6.09E-06) and “filamentous actin” (P = 7.43E-06), which play an active role in wound healing and tissue regeneration (24, 25) (**Supplementary Table 10**). Although DNA methylation at only one site (cg26426898 on chromosome 20, annotated to *SCRT2* and *C20orf54*) was identified as significantly different (P < 2.0E-07) at both POD1 and POD4/7 compared to BL, there was a strong correlation of changes observed at POD1 and POD4/7 for DMPs identified at both time-points (POD1 DMPs: corr = 0.925, P = 2.86E-05, **Figure 2C**; POD4/7 DMPs: corr = 0.745, P = 6.00E-09, **Figure 2D**) indicating that surgery-induced DNA methylation changes are relatively stable until the time of discharge.

**Table 3.**
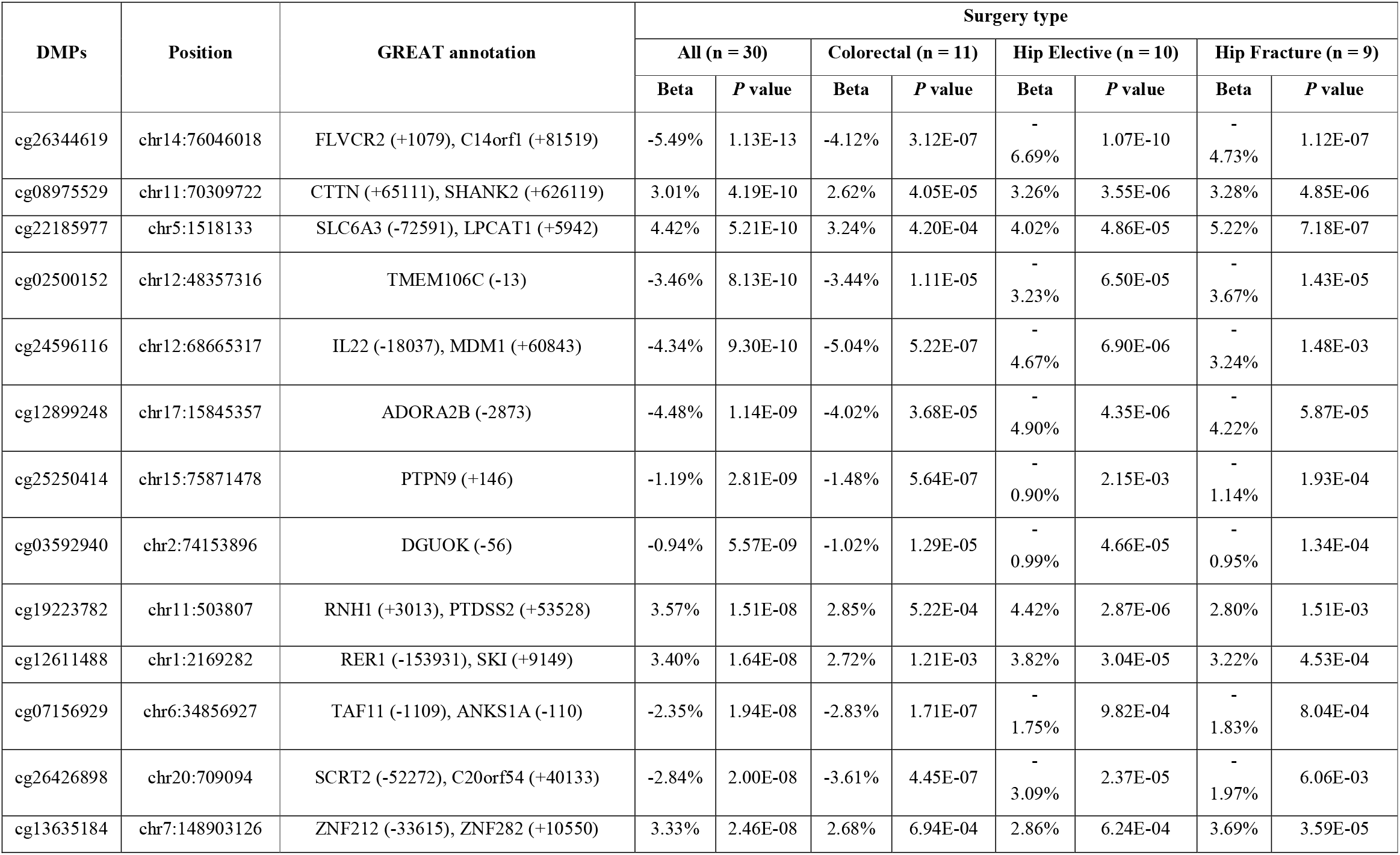
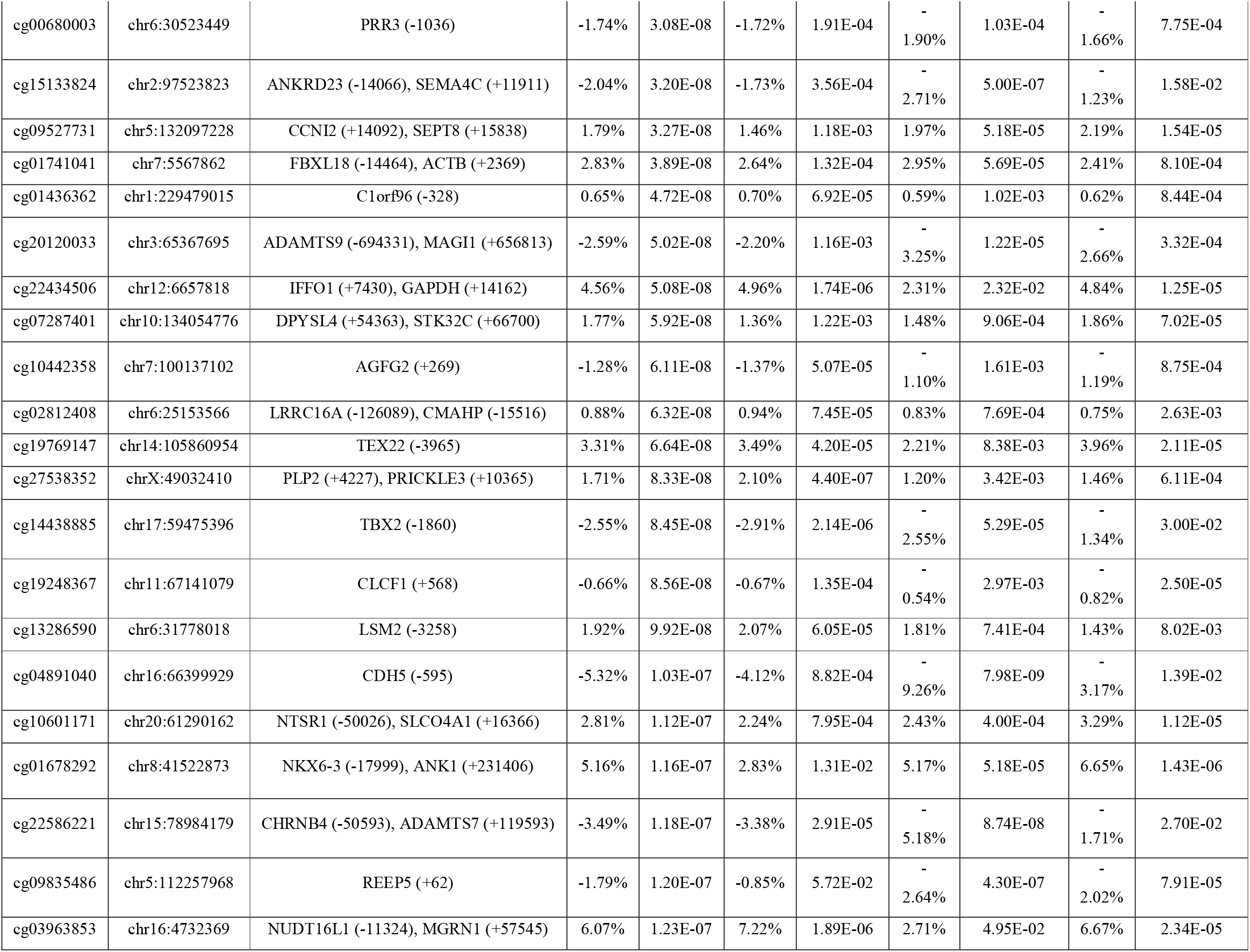
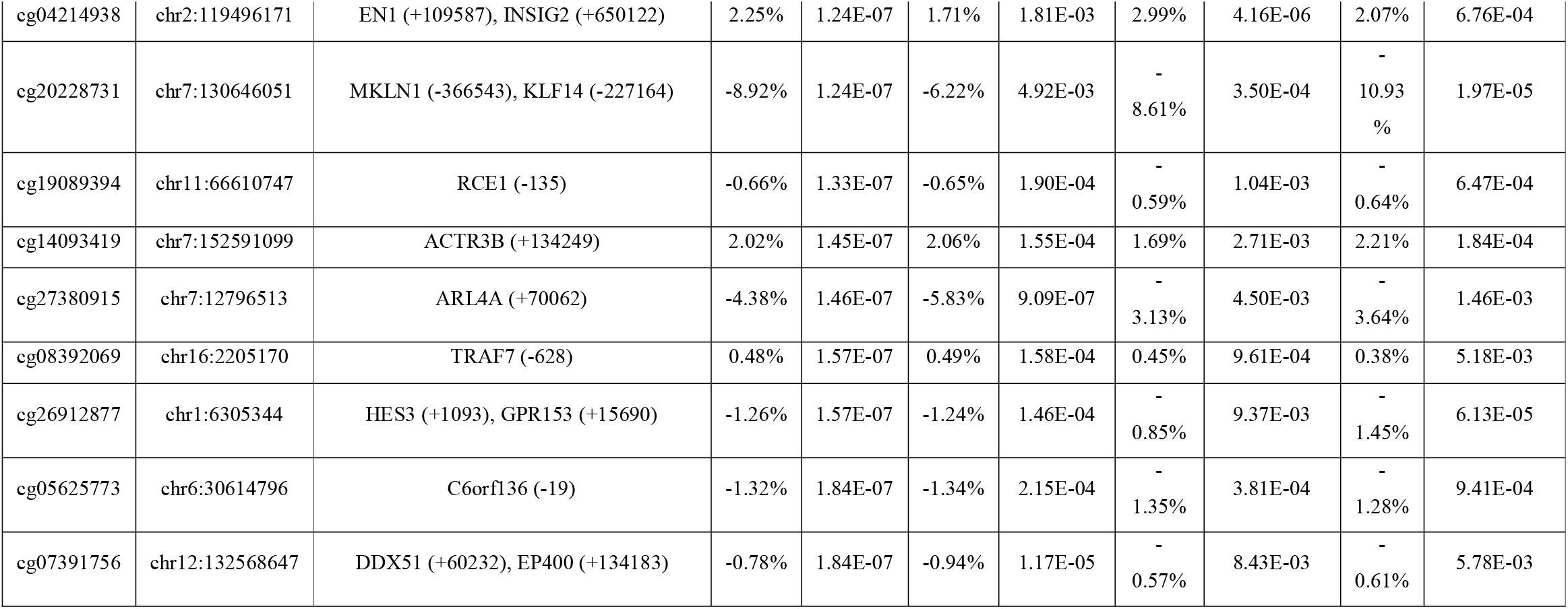
Sites characterized by significant changes in DNA methylation on post-operative day 4 to 7 compared to baseline. Shown for each DMP are the genomic location (hg19) and genic annotation derived using GREAT (35). Association statistics are presented across all patients, and separately for each surgery type.

### Surgery-associated DNA methylation differences identified by the Illumina 450K were robustly validated using bisulfite-pyrosequencing

We used bisulfite-pyrosequencing to confirm surgery-associated changes in DNA methylation at several DMPs, validating the results of our Illumina 450K analyses and finding highly consistent results across platforms. First, we re-quantified DNA methylation at cg24501381, one of the top-ranked DMPs from our initial analysis uncorrected for cell-types (average DNA methylation change = −4.99%, P = 5.69E-09) that is annotated to *CCDC30* on chromosome 1. There was a highly significant correlation between Illumina 450K array and bisulfite-pyrosequencing datasets (P = 1.04E-23, **Supplementary Figure 10**), with the pyrosequencing data confirming the surgery-associated reduction in DNA methylation (average DNA methylation change = −4.37%, P = 8.75E-06, **Supplementary Figure 11**). Second, we re-quantified DNA methylation at cg26344619, the top-ranked DMP at POD4/7, again finding a highly significant correlation between 450K array and bisulfite-pyrosequencing datasets (P = 5.38E-19, **Supplementary Figure 12**), with the pyrosequencing data confirming the surgery-associated reduction in DNA methylation (average DNA methylation change = −4.90%, P = 7.07E-05, **Supplementary Figure 13**).

### Heterogeneity in DNA methylation changes between different types of surgery

The DNA methylation changes at POD1 and POD4/7 described above are relatively consistent across the three types of major surgery (colorectal elective surgery, hip elective surgery, and hip fracture surgery) (**Figure 2A** and **Figure 2B**). It is, however, possible that there will be unique effects specific to individual surgery-type groups or between emergency and elective surgery. The three groups were relatively well-matched for most demographic and surgical parameters; although there were no differences in smoking status, marital status, Charlson’s comorbidity score, Katz index, ASA classification, or blood loss between surgery types, patients undergoing colorectal surgery were younger (P = 1.24E-02) and were exposed to a longer duration of surgery (P = 1.13E-02) than the other groups (**Table 1**). Anaesthesia time was also longer in patients in the colorectal surgery group than those undergoing hip elective surgery (P = 1.62E-03). We used an analysis model designed to identify changes in DNA methylation that differed between surgery types (see **Methods**), finding eight surgerytype-specific DMPs (P < 2.0E-07) at POD1 (**Supplementary Table 11** and **Supplementary Figure 14**) and eleven at POD4/7 (**Supplementary Table 12** and **Supplementary Figure 15**).

## Discussion

In this study we characterized longitudinal changes in DNA methylation in purified peripheral blood PBMCs collected from elderly patients undergoing major surgery. Our primary analyses focused on comparing samples collected at baseline to those collected immediately post-operatively and at discharge from hospital. We observed rapid changes in DNA methylation following surgery, with widespread differences becoming manifest before the morning of postoperative day1. The acute changes in DNA methylation induced by surgery are relatively stable in the post-operative period, generally persisting until discharge from hospital.

We found that major surgery is associated with acute changes in DNA methylation at sites annotated to immune system genes, reflecting the observed changes in serum-levels of markers associated with surgical trauma, including C-reactive protein (CRP) and interleukin 6 (IL-6), in the same samples. This physiological response involves dynamic changes in gene regulation, and the acute shifts in DNA methylation observed here parallel the immunological responses to major surgery that underlie the post-operative recovery (7). It is therefore plausible that in addition to modulating the immune response to major surgery, DNA methylation might be involved in regulating susceptibility to certain postoperative complications such as nosocomial infection, delirium, and cognitive dysfunction (4, 5). Given the growing need to optimise surgical recovery in our aging population, our findings highlight how perioperative molecular approaches might provide insight into the pathophysiology of surgical complications, and future work will focus on stratifying individuals to identify methylomic signatures of poor outcome (26). Although many of the observed changes in DNA methylation are consistent across the three types of surgery, there is notable heterogeneity between surgery types at certain loci. These differences may reflect different types of surgical procedures, emergency and elective surgery, or even the tissues operated upon, the surgical access type and underlying pathology.

This study has several strengths and weaknesses that should be considered when interpreting the results our analyses. First, we implemented a prospective study design, longitudinally sampling individuals before and at several time-points after major surgery. The analysis of sequential samples from the same individual that negates many of the important confounds that can influence molecular epidemiology as we are using each patient as their own baseline compared to an external reference level from a heterogeneous population and we can control for variables such as age, sex and exposure (e.g. to smoking and medication) (27). Second, although we isolated PBMCs from fresh whole blood, cellular heterogeneity remains an important potential confounder especially given the known blood-cell changes known to occur following surgery. Of note, we were able to correct for blood cell proportions, albeit only for major cell-types, and show that our derived estimates reflected directly measured cell-type levels. Our results support the usefulness of cell-correction algorithms in epigenetic epidemiology and highlight the importance of adjusting for cell type composition for the epigenetic research in perioperative period. Third, we quantified DNA methylation using the Illumina 450K array; although this platform interrogates sites annotated to the majority of genes, the actual proportion of sites across the genome interrogated by this technology is relatively low, with a predominant focus on CpG-rich promoter regions. It will be important for future studies to explore surgery-induced changes in DNA methylation across regions not well-covered by the Illumina 450K array. Of note, we used bisulfite-Pyrosequencing to validate surgery-induced DNA methylation changes at several loci, confirming the robustness of the 450K array for identifying real DNA methylation changes. Finally, the inclusion of patients undergoing multiple surgery types enabled us to explore the consistency of DNA methylation changes following major surgery. Despite this, however, individual surgery-type groups were relatively small and future work is needed to further interrogate genomic changes specific to each group. Surgery is a complex procedure and involves a large combination of factors that could influence DNA methylation including the surgical approach (open versus minimally-invasive), differences in anaesthetic techniques used, differences in the tissues undergoing surgery, and different rates of complications and compounding factors. Future work in larger surgical cohorts will enable us to address these factors.

## Conclusion

To our knowledge, this represents the first study to systematically explore acute changes in gene regulation following major surgery. We identified acute surgery-associated changes in DNA methylation which are relatively stable until discharge from hospital. Taken together, our results highlight the dramatic alterations in gene regulation induced by invasive surgery, primarily reflecting upregulation of the immune system in response to trauma.

## Methods

### Participants and sample collection

Potential participants were identified by clinicians during routine clinical practice. Between July 2014 and January 2015, a total of 55 patients undergoing routine/emergency surgery at the Royal Devon & Exeter Hospital (Exeter, UK) were recruited via the Royal Devon and Exeter Tissue Bank (RDETB). The ethically approved RDETB (REC ref: 11/SW/0018) was set up to proactively collect and store tissue available from routine clinical procedures for studies examining disease specific biomarkers and is facilitated through the NIHR Exeter Clinical Research Facility (ECRF) https://exetercrfhihr.org/about/rde-tissue-bank/. Patients with known dementia, cognitive decline (defined as < 9/10 points on the Abbreviated Mental Test (28)), delirium before surgery, early discharge within 3 days after surgery were excluded from participating in the study. Written informed consent was obtained from all participants. Pre-operatively, patients were assessed using the Katz index (29), the Charlson’s comorbidity score (30) and the American Society of Anesthesiologists classification, with the absence of delirium was assessed using the Confusion Assessment Method (31) were assessed. Peripheral whole blood samples were obtained at three time points: 1) immediately before surgery at baseline (“BL”), 2) in the morning on post-operation day 1 (“POD1”) and 3) on day 4-7 immediately prior to discharge (“POD4/7”). Whole blood was taken by a trained research nurse in addition to clinical examination and samples were immediately transferred to the research team for isolation of peripheral blood mononuclear cells (PBMCs). Out of the 55 patients recruited to the study, a complete set of biological and clinical samples were collected from 30 patients. A number of patients were excluded; unstable / severe clinical condition (n = 2), cancelled surgery (n = 4), failed sample collection (n = 2), history of delirium (n = 2), or early discharge and a failure to collect three blood samples (n=15). An overview of the patients included in our final analysis dataset is given in **Supplementary Table 1**.

### IL-6, CRP, and blood component measurements

Serum was collected from fresh blood samples by centrifugation at 1,600 x g at room temperature for 20 minutes and stored at −80°C until use. Serum IL-6 levels were measured using the Quantikine^®^ Enzyme Linked Immuno-Sorbent Assay Human IL-6 Kit (R&D Systems) following the manufacturers standard protocol. Each experiment was performed in duplicate and calibrated using standards. Serum levels of CRP and other blood components were measured in the hospital clinical chemistry laboratory. Laboratory-derived data measured within 3 days before surgery were accepted as baseline.

### Genome-wide DNA methylation profiling

PBMCs were isolated from fresh whole blood, 15 minutes to 2 hours after collection, using Mononuclear Cell Preparation Tubes (BD, Vacutainer^®^ CPT). Ficoll (BD) separated PBMCs were centrifuged at 1,600 x g at room temperature for 20 minutes, and then washed twice with 5ml PBS. Isolated PBMCs were resuspended in RNAprotect cell reagent (Qiagen) and stored at −80 °C. Genomic DNA was extracted using the AllPrep^®^ DNA/RNA minikit (Qiagen, Redwood City, CA) and bisulfite converted using the EZ-96 DNA Methylation-Gold kit (Zymo research, Irvine, CA) following the manufacture’s standard protocol. Aliquoted bisulfite converted DNA was amplified with bisulfite specific primers to confirm conversion efficiency prior to array analysis. Genome-wide DNA methylation was profiled using the Illumina Infinium HumanMethylation450 BeadChip (Illumina Inc. San Diego, CA) (“450K array”) and scanned on an Illumina iScan. Following stringent quality control (QC) of the raw data, the data was normalized using the *dasen* function in the *wateRmelon* package for a combination of background adjustment and between-array quantile normalization (32). Probes with non-specific sequences and those containing common (Minor allele frequency > 5%) single nucleotide polymorphism probes in European population were removed (33) leaving 427,353 probes for analysis.

### Bisulfite-pyrosequencing

Bisulfite-pyrosequencing was performed to validate DNA methylation data from the Illumina 450K array for two CpG sites using specific primers (**Supplementary Table 13**) designed using PyroMark Assay Design software 2.0 (Qiagen). DNA methylation was quantified from intensities captured in the PyroMark Q24 pyrosequencer (Qiagen) based on duplicated bisulfite polymerase chain reaction amplification.

### Statistical analyses

All statistical analyses were performed in the R statistical environment (version, 3.3.2). Surgery-induced DNA methylation across the genome was first examined using paired t-test for 427,353 DNA methylation sites included in our dataset following QC. We then used a multi-level linear regression model fitted for each DNA methylation site with fixed effects for estimated cell counts (derived using the Epigenetic Clock Software (20) : https://labs.genetics.ucla.edu/horvath/dnamage/), age, sex, and processing batch and a random effect for individual using the *lme4* R package (34). The EWAS of C-reactive protein levels were performed using the same package with fixed effects for blood collection point, surgery type, estimated cell counts, age, sex, and processing batch. For Illumina array analyses we considered used a significance threshold of P < 2.0E-7 for genome wide significance and P < 5.0E-05 to select probes for GO pathway analyses. Demographic data were tested using analysis of variance (ANOVA) or a Fisher’s exact test. Laboratory-derived blood data alterations following surgery were tested using a paired t-test.

## Abbreviations

ANOVA: analysis of variance
BL: baseline
CpG: cytosine-guanine dinucelotide
CRP: C-reactive protein
CRF: clinical research facility
DMP: differentially methylated position
DMR: differentially methylated region
DNA: deoxyribonucleic acid
ECRF: Exeter Clinical Research Facility
EWAS: epigenome-wide association study
GO: gene ontology
IL-6: Interleukin 6
PBMC: peripheral blood mononuclear cells
POD: post-operative day
QC: quality control
RDETB: Royal Devon and Exeter Tissue Bank
TSS: transcription start site

## Declarations

### Ethics approval and consent to participate

This study was approved by the RDETB Steering Committee under the terms of the overall RDETB ethical approval (REC ref: 11/SW/0018). All participants provided written informed consent for participation.

### Consent for publication

Not applicable.

### Availability of data and materials

Raw data and analysis scripts are available from the authors on request.

### Authors contributions

J.M., N.S. and R.S. conceived the study and managed the project. B.K. and R.S. coordinated clinical sampling. B.K., J.C., N.S. and I.D. recruited patients for the study. J.B. and B.C. undertook DNA methylation profiling. R.S. performed primary bioinformatics analyses. E.H. and O.K. provided statistical and bioinformatics support. F.J. undertook IL-6 ELISA analysis. J.M. and R.S. drafted the manuscript. All authors read and approved the final manuscript prior to submission.

## Acknowledgements

This project was supported by the National Institute for Health Research (NIHR) Exeter Clinical Research Facility (Exeter CRF). The views expressed are those of the author(s) and not necessarily those of the NIHR or the Department of Health and Social Care. The NIHR Exeter CRF is a partnership between the University of Exeter Medical School College of Medicine and Health, and Royal Devon and Exeter NHS Foundation Trust.

