## Supplementary Figures for "Major surgery induces acute changes in DNA methylation associated with activation of the immune response"

**Supplementary Figure 1 – An overview of the study design.** We recruited a cohort of 30 elderly patients (average age = 77.9 years) undergoing major surgery (colorectal elective surgery (n = 11), elective hip replacement surgery (n = 10), and hip fracture surgery (n = 9)). From each individual, detailed blood measures were collected and DNA methylation was profiled using the Illumina HumanMethylation450 array in DNA samples isolated from peripheral blood mononuclear cells (PBMCs) collected at three time-points: i) immediately before surgery (baseline), ii) in the morning of post-operative day 1 (POD1) and iii) on post-operative day 4 to 7 (POD4/7) before discharge from hospital.

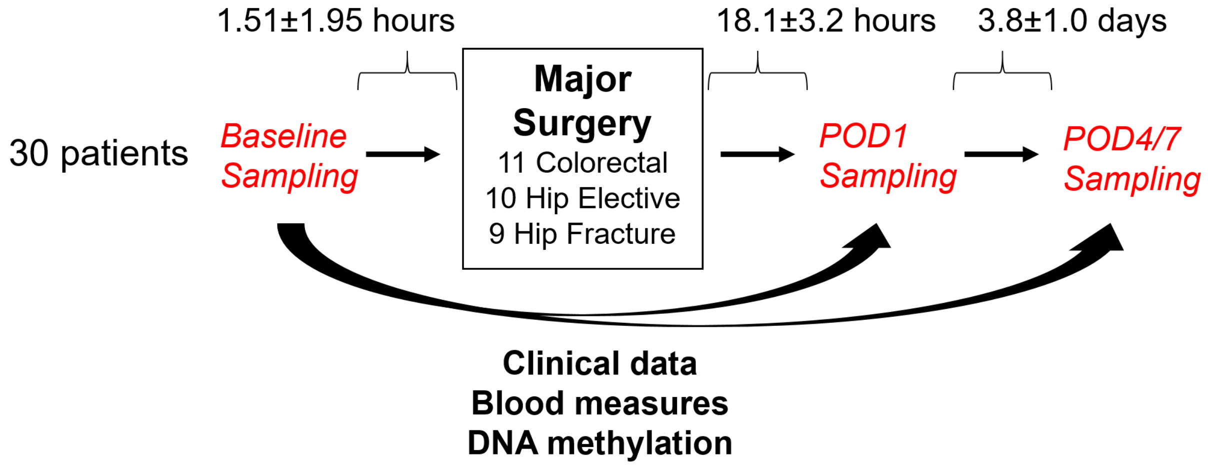

**Supplementary Figure 2 – Manhattan plot for acute DNA methylation changes associated with major surgery.** Within each individual, DNA methylation was compared between BL and POD1. Red line = experiment-wide significance threshold (P < 2.0E-07). Blue line = discovery significance threshold (P < 5.0E-05) used for gene ontology analysis.

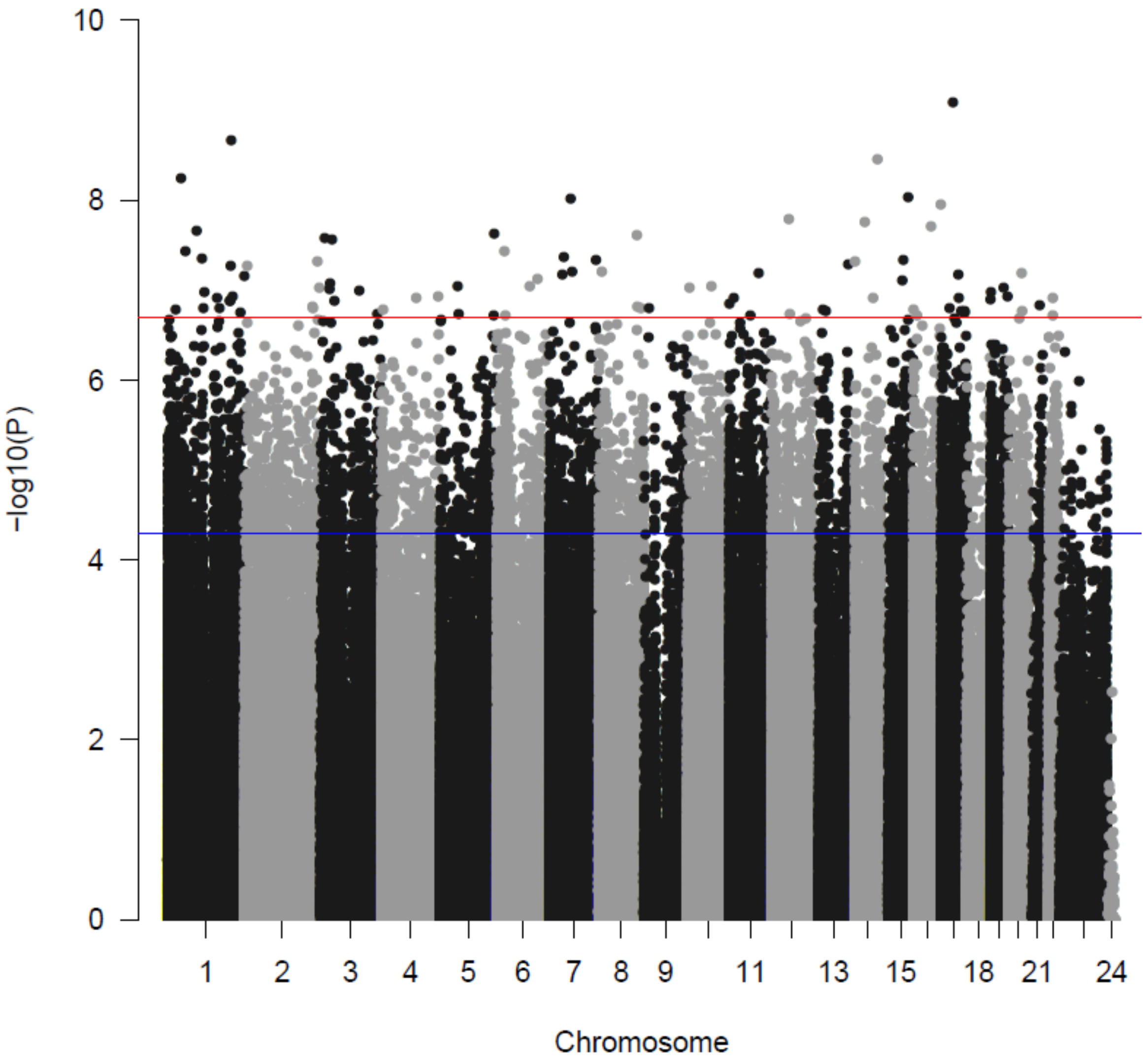

**Supplementary Figure 3 – Surgery is associated with acute changes in multiple blood plasma measures.** Significant acute changes were observed in plasma-based blood measures including an increase in bilirubin (baseline: mean bilirubin = 8.2 μmol/l; POD1: mean bilirubin = 13.3 μmol/l, P = 4.19E-03), and decreases in alkaline phosphatase (baseline: mean alkaline phosphatase = 82.8 iu/l; POD1: mean alkaline phosphatase = 66.9 iu/l, P = 1.15E-05), albumin (baseline: mean albumin = 44.2 g/l; POD1: mean albumin = 35.6 g/l, P = 1.19E-10), and sodium (baseline: mean sodium = 137.9 mEq/l; POD1: mean sodium = 135.7 mEq/l, P = 2.41E-04), haemoglobin (baseline: mean haemoglobin = 123.4 g/l; POD1: mean haemoglobin = 102.7 g/l, P = 5.33E-09) and haematocrit (baseline: mean haematocrit = 36.9 %; POD1: mean haematocrit = 30.6 %, P = 6.16E-09). Error bar shows the 95% confidence interval.

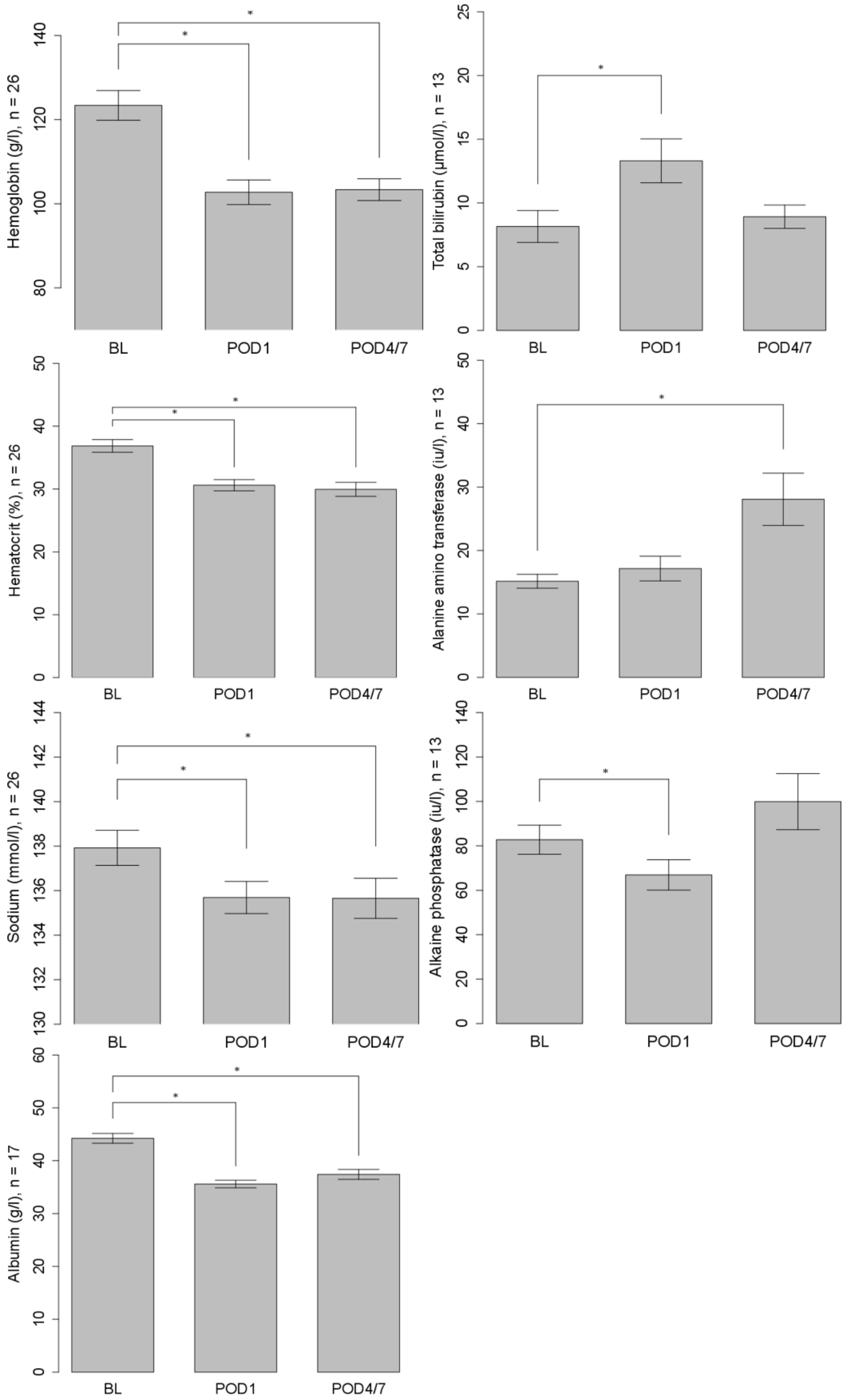

**Supplementary Figure 4 - Surgery is associated with significant changes in blood cellular composition.** Significant acute changes were observed in blood cellular composition including an increase in white blood cell count (baseline: mean white blood cell count = 9.73 *1.000/µl; POD1: mean white blood cell count = 11.35 *1.000/µl, P = 4.08E-02), neutrophil count (baseline: mean neutrophil count = 7.57 *1.000/µl; POD1: mean neutrophil count = 9.33 *1.000/µl, P = 3.00E-02) , and monocyte count (baseline: mean monocyte count = 0.68 *1.000/µl; POD1: mean monocyte count = 0.85 *1.000/µl, P = 5.64E-03), and decreases in red blood cell count (baseline: mean red blood cell count = 4.04 *10E6/µl; POD1: mean red blood cell count = 3.38 *10E6/µl, P = 3.24E-09), platelets (baseline: mean platelets = 247.5 *1.000/µl; POD1: mean platelets = 212.5 *1.000/µl, P = 8.66E-07), lymphocyte count (baseline: mean lymphocyte count = 1.28 *1.000/µl; POD1: mean lymphocyte count = 1.11 *1.000/µl, P = 2.34E-02), eosinophil count (baseline: mean eosinophil count = 0.15 *1.000/µl; POD1: mean eosinophil count = 0.04 *1.000/µl, P = 4.03E-03) and basophil count (baseline: mean basophil count = 0.05 *1.000/µl; POD1: mean basophil count = 0.02 *1.000/µl, P = 2.08E-03). Error bar shows the 95% confidence interval.

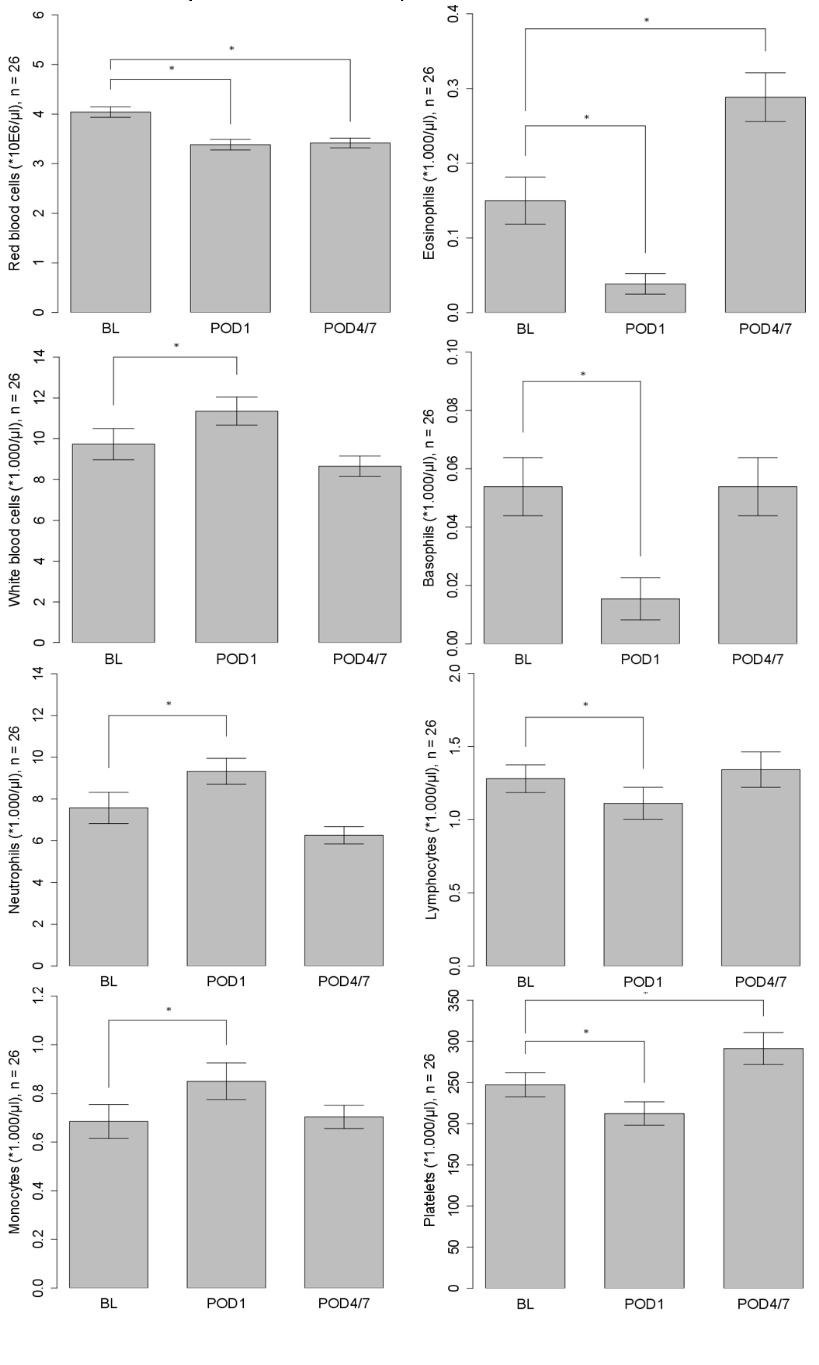

**Supplementary Figure 5 – Blood cell proportion estimates derived from DNA methylation data are correlated with empirical cell-type measures.** Shown is the correlation between empirically-measured ratios between monocytes and lymphocytes and those derived from DNA methylation data in peripheral mononuclear cells (n = 86, r = 0.713, *P* = 8.53E-15).

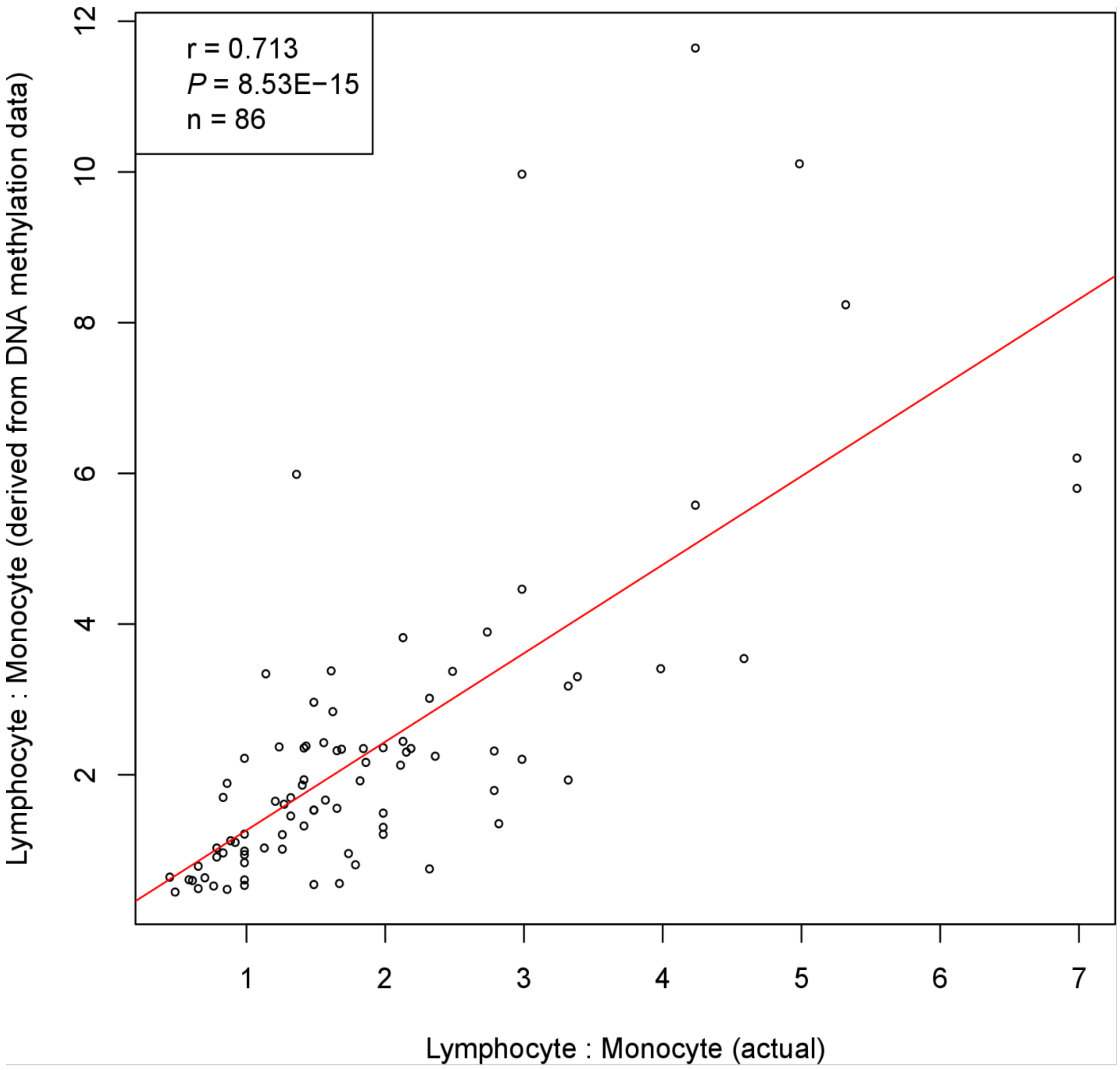

**Supplementary Figure 6 - There was a strong correlation for effect sizes (r = 0.955, P = 2.20E-47) between models (corrected and uncorrected for derived cell proportions) for DMPs identified in our uncorrected model.**

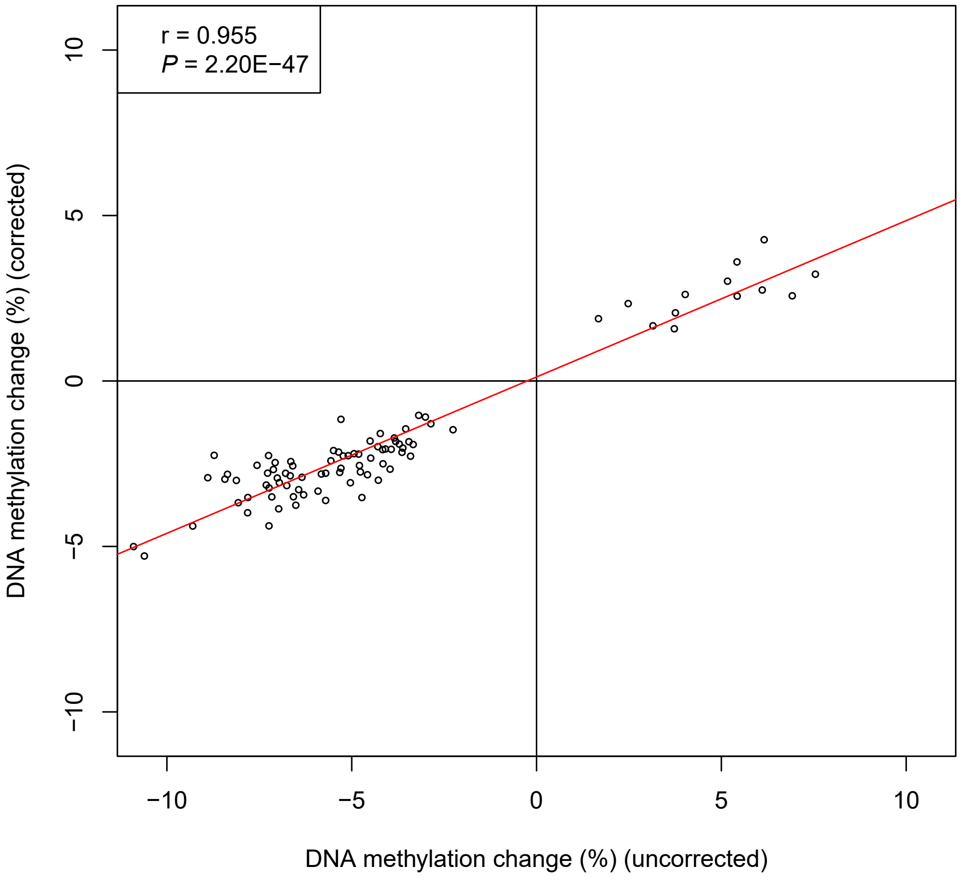

**Supplementary Figure 7 – There was a strong correlation for P values (r = 0.621, P = 6.45E-11) between models (corrected and uncorrected for derived cell proportions) for DMPs identified in our model uncorrected for blood cell-type proportions.**

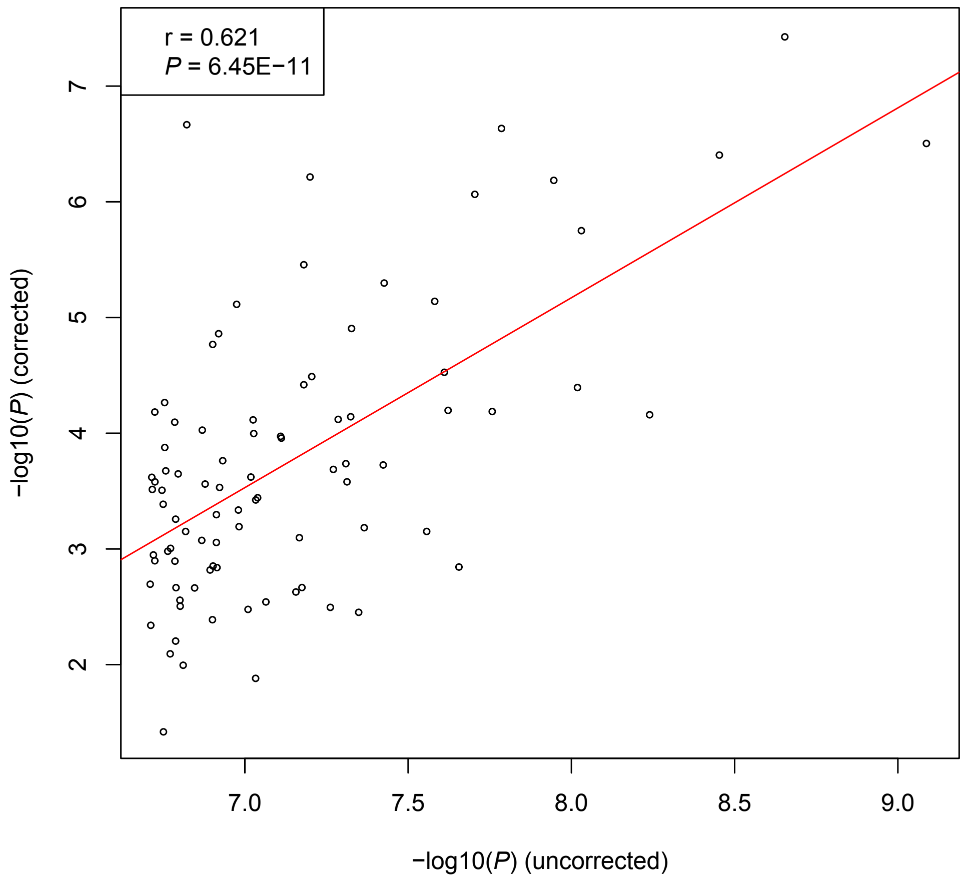

**Supplementary Figure 8 – Manhattan plot for acute DNA methylation changes associated with major surgery in a model correcting for blood cell-type proportions.** Within each individual, DNA methylation was compared between BL and POD1. Red line = experiment-wide significance threshold (P < 2.0E-07). Blue line = discovery significance threshold (P < 5.0E-05) used for gene ontology analysis.

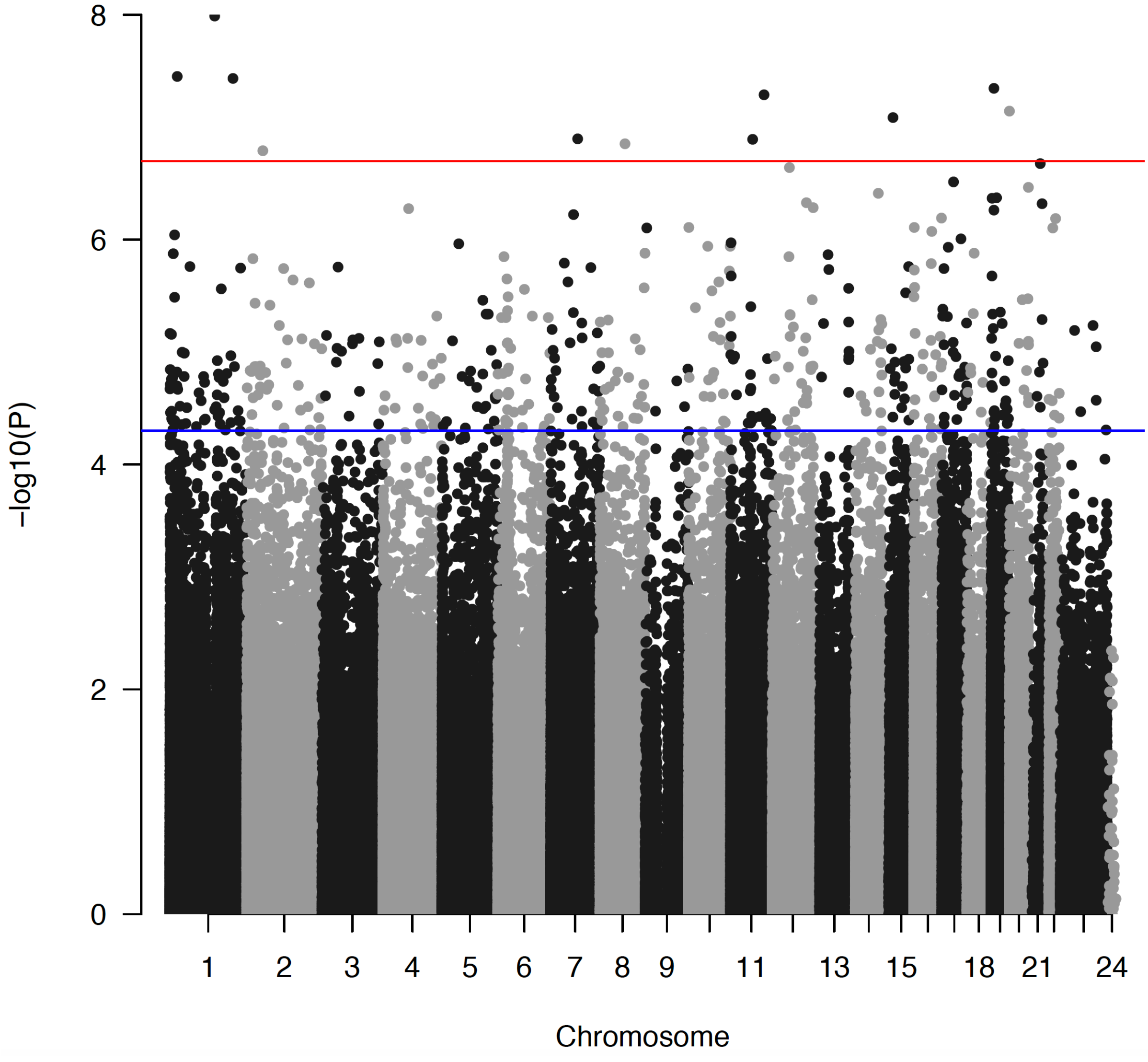

**Supplementary Figure 9 - Manhattan plot for DNA methylation changes between BL and POD4/7 associated with major surgery in a model correcting for blood cell-type proportions.** Red line = experiment-wide significance threshold (P < 2.0E-07). Blue line = discovery significance threshold (P < 5.0E-05) used for gene ontology analysis.

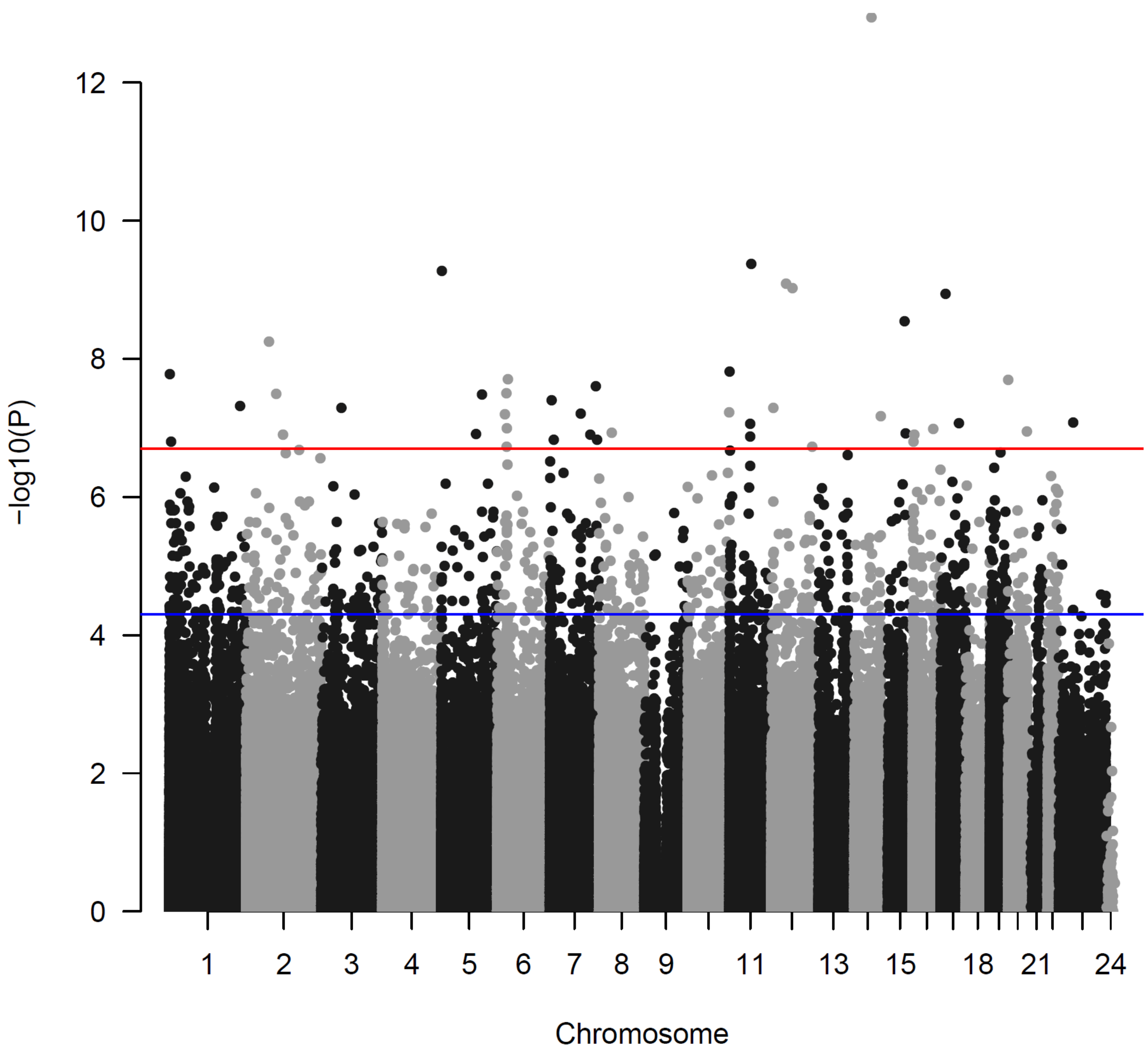

**Supplementary Figure 10 – Comparison of bisulfite-pyrosequencing and Illumina 450K array data for DNA methylation at cg24501381.** There was a highly significant correlation between 450K array and bisulfite-pyrosequencing datasets (P = 1.04E−23).

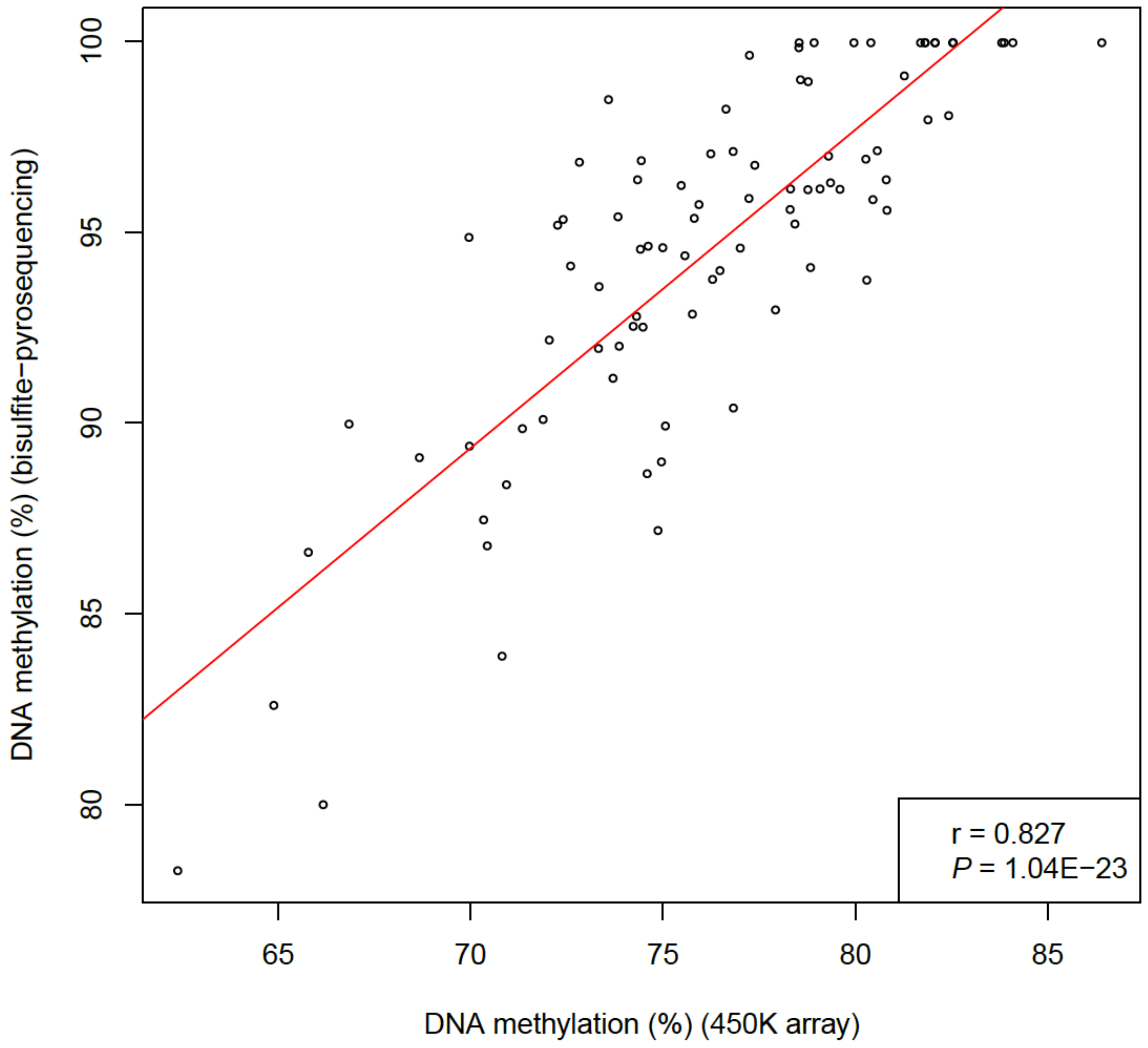

**Supplementary Figure 11 – Bisulfite-pyrosequencing data confirmed a significant surgery-associated reduction in DNA methylation at cg24501381.**

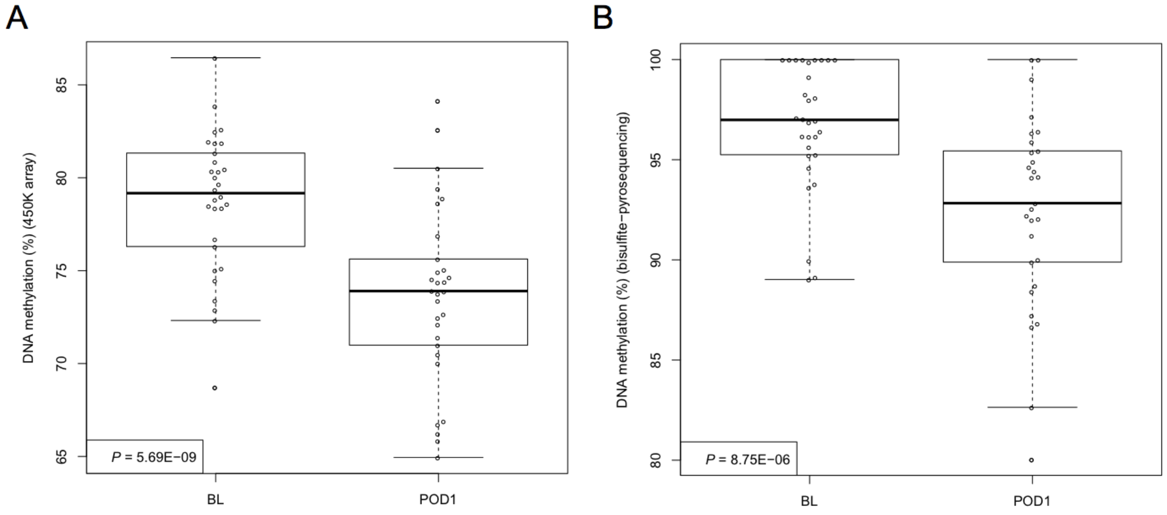

**Supplementary Figure 12 – Comparison of bisulfite-pyrosequencing and Illumina 450K array data for DNA methylation at cg26344619.** There was a highly significant correlation between 450K array and bisulfite-pyrosequencing datasets (P = 5.38E-19).

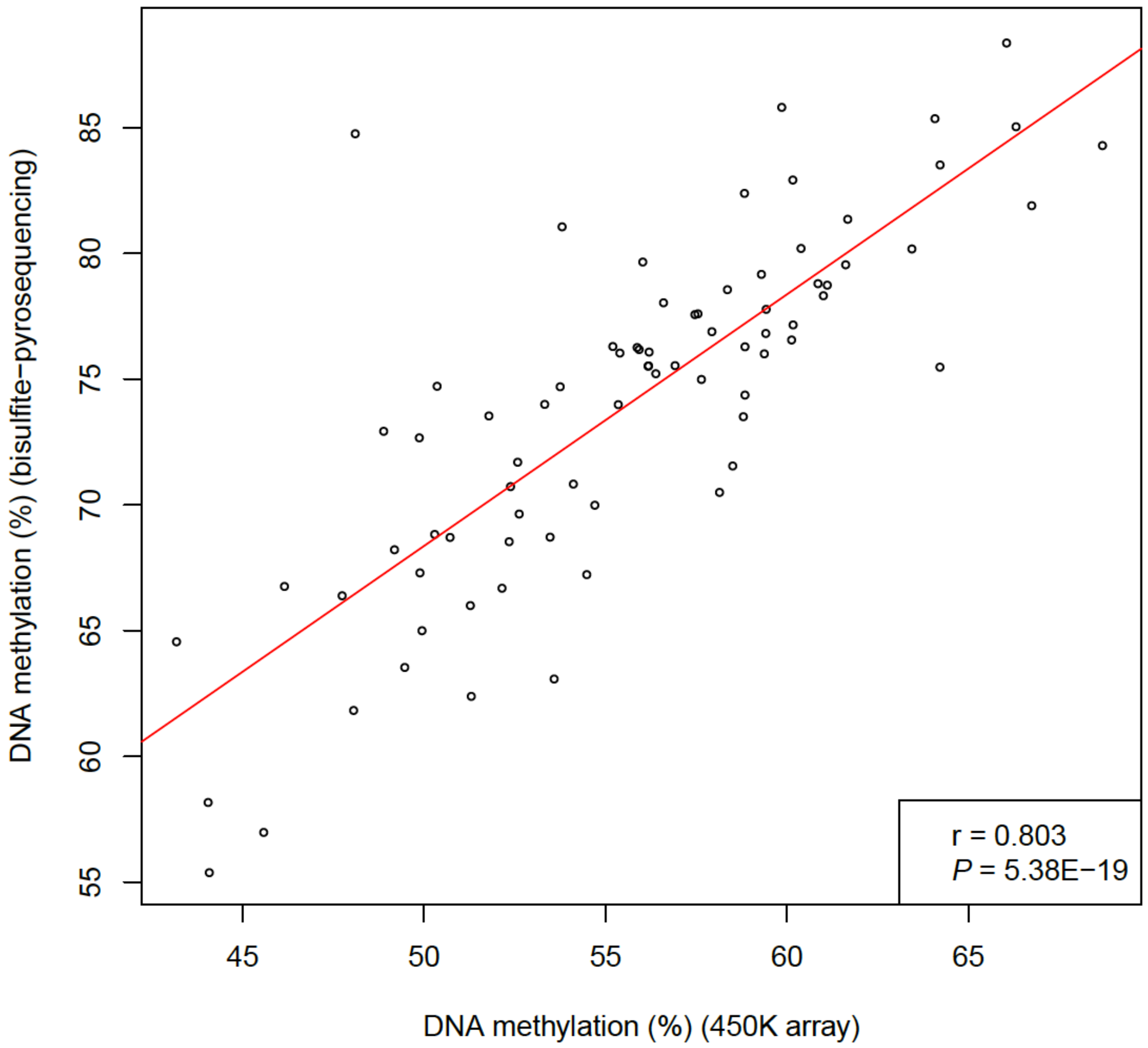

**Supplementary Figure 13 – Bisulfite-pyrosequencing data confirmed a significant surgery-associated reduction in DNA methylation at cg26344619.**

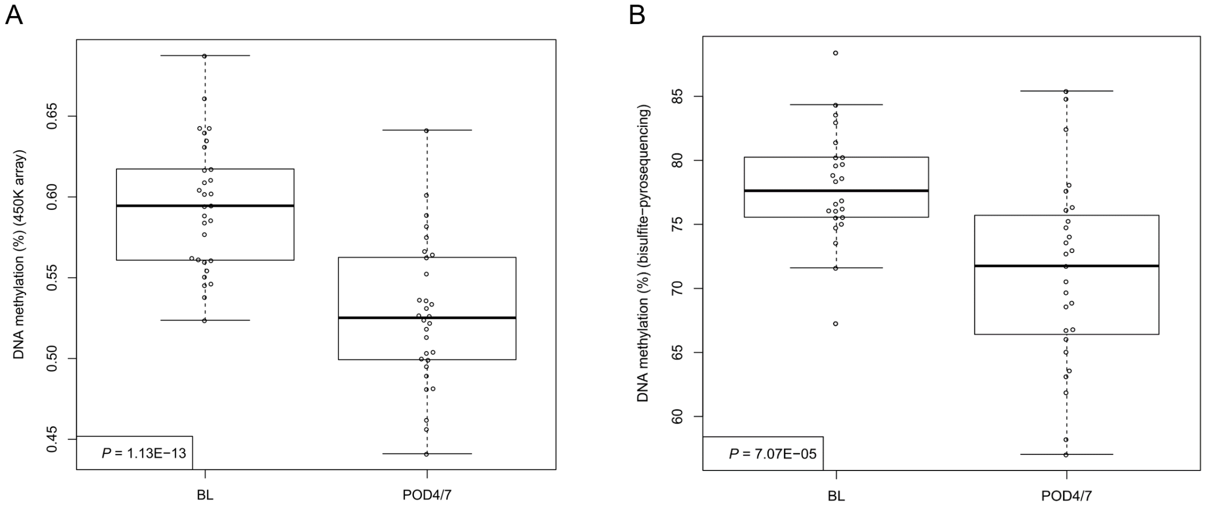

**Supplementary Figure 14 - Surgery-type-specific DMPs (P < 2.0E-07) at POD1.** Data for each of these eight DMPs is given in **Supplementary Table 11**.

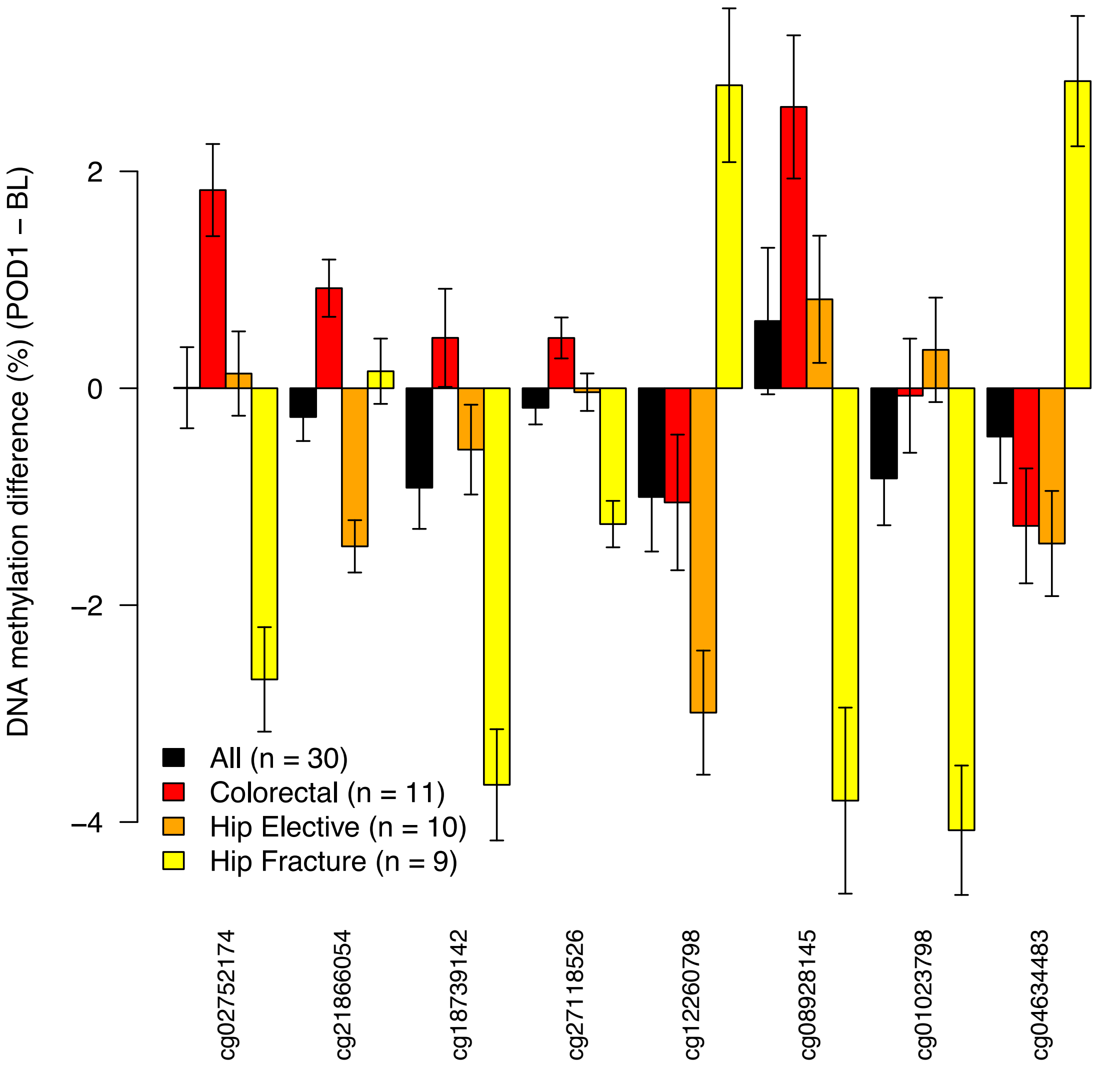

**Supplementary Figure 15 - Surgery-type-specific DMPs (P < 2.0E-07) at POD4/7.** Data for each of these eleven DMPs is given in **Supplementary Table 12**.

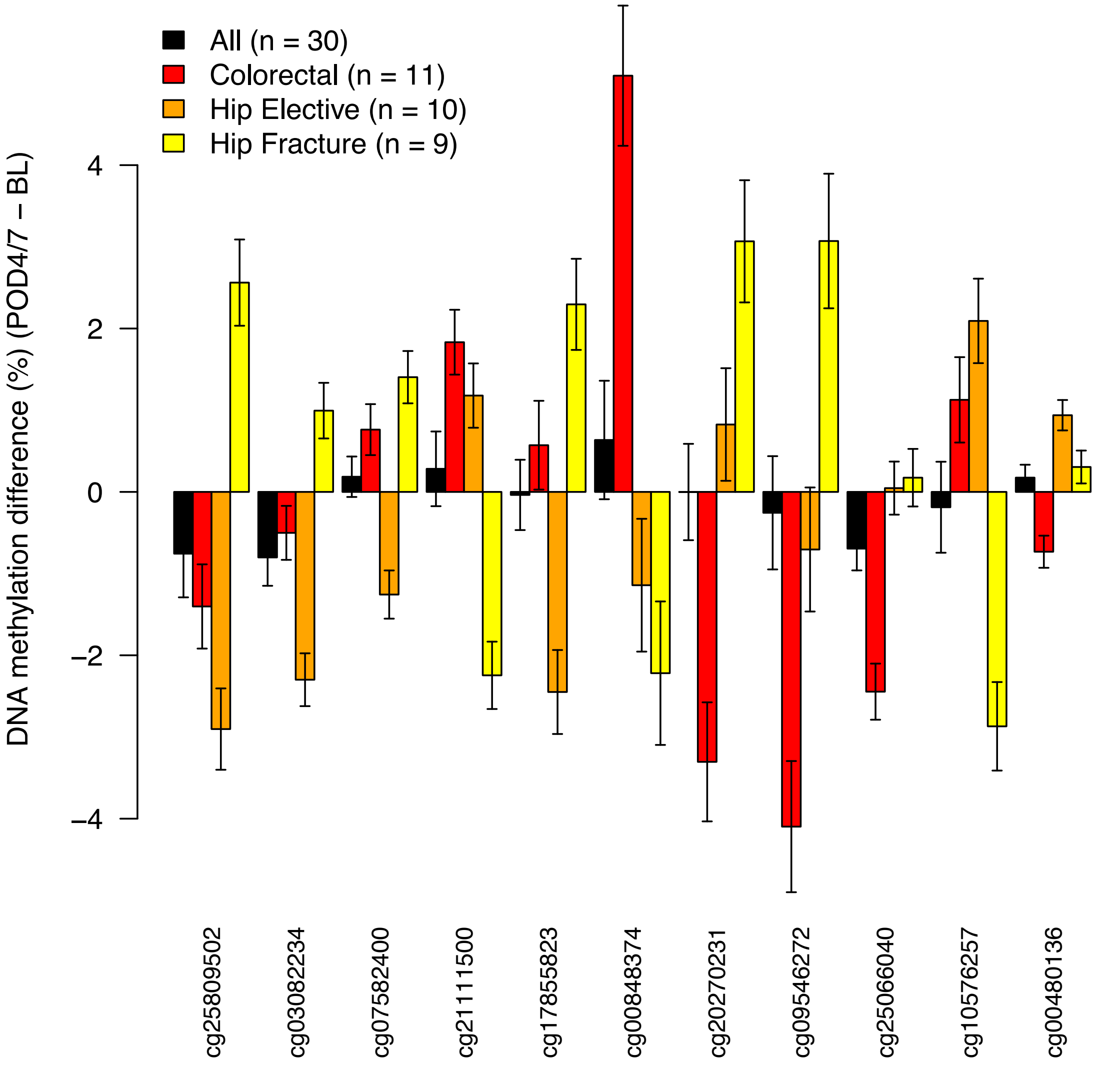
